# BAP1 deubiquitinase is a potent repressor of fetal hemoglobin biosynthesis

**DOI:** 10.1101/346619

**Authors:** Lei Yu, Natee Jearawiriyapaisarn, Mary P. Lee, Tomonori Hosoya, Qingqing Wu, Greggory Myers, Kim-Chew Lim, Ryo Kurita, Yukio Nakamura, Anne B. Vojtek, Jean-François Rual, James Douglas Engel

## Abstract

Human globin gene production transcriptionally “switches” from fetal to adult synthesis shortly after birth, and is controlled by macromolecular complexes that enhance or suppress transcription by cis-elements scattered throughout the locus. The DRED repressor is recruited to the ε- and γ-globin promoters by the orphan nuclear receptors TR2 (NR2C1) and TR4 (NR2C2) to engender their silencing in adult erythroid cells. Here we found that nuclear receptor corepressor-1 (NCoR1) is a critical component of DRED that acts as a scaffold to unite the DNA binding and epigenetic enzyme components (e.g. DNMT1 and LSD1) that elicit DRED function. We also describe a potent new regulator of γ-globin repression: the deubiquitinase BAP1 is a component of the repressor complex whose activity maintains NCoR1 at sites in the β-globin locus, and BAP1 inhibition in erythroid cells massively induces γ-globin synthesis. These data provide new mechanistic insights through the discovery of novel epigenetic enzymes that mediate γ-globin gene repression.

## Introduction

The fetal to adult developmental switch in the human β-globin gene locus involves the alteration in transcription from synthesis of the two nearly identical fetal γ-globin genes to the adult β-globin gene, a process which is elicited through the cumulative activities of transcriptional activators and repressors that bind to their cognate cis-regulatory elements in the β-globin locus (Bulger and Groudine, 1999; Engel and Tanimoto, 2000).

Previously, we identified a multi-subunit transcriptional repressor complex that we named DRED (Direct Repeat Erythroid-Definitive). DRED binds with high affinity to direct repeat (DR1) elements located in the human embryonic ε-globin and fetal γ-globin, but not the adult β-globin, promoters and it represses γ-globin transcription in definitive adult erythroid cells (Tanabe et al., 2002; Tanabe et al., 2007). Based on the results of protein affinity purification followed by mass spectrometric analysis, we proposed that the “core” DRED complex was a tetramer composed of a heterodimer of the orphan nuclear receptors TR2 (NR2C1) and TR4 (NR2C2), which bind directly to the DR1 elements in the ε- and γ-globin gene promoters, and the two corepressor enzymes DNA methyltransferase 1 (DNMT1) and lysine-specific demethylase 1 (LSD1 or KDM1a)(Cui et al., 2011).

We and others have shown that pharmacological inhibition of the enzymatic activities of these DRED corepressor enzymes, DNMT1 (with 5-azacytidine or decitabine) and LSD1 (with tranylcypromine or RN-1), results in the induction of fetal γ-globin synthesis in adult definitive erythroid cells (Clegg et al., 1983; Cui et al., 2015a; Cui et al., 2015b; DeSimone et al., 1982; Ley et al., 1983; McCaffrey et al., 1997; Molokie et al., 2017; Rivers et al., 2016; Rivers et al., 2015; Shi et al., 2013). These inhibitory strategies were proposed as possible pathways that could lead to robust fetal hemoglobin (HbF) induction, and therefore a potentially effective therapeutic strategy to treat the β-globinopathies (sickle cell anemia and β-thalassemia)(Suzuki et al., 2014). However, all of the corepressor enzymes identified to date are widely or ubiquitously expressed and are therefore certain to participate in other biological functions. Therefore, their potential as therapeutic agents for HbF induction may be dependent on their tissue abundance and exposure to the bloodstream.

Hydroxyurea is the current treatment standard for sickle cell disease, but has significant associated problems, and no other treatment options are currently available that might achieve sufficient levels of HbF induction that would ameliorate both the symptoms and pathophysiology caused by red blood cell sickling (Ngo et al., 2012; Noguchi et al., 1988; Wood et al., 1976). Similarly, there is no current pharmacological treatment for patients with β^major^-thalassemia that would improve imbalanced hemoglobin chain synthesis. Thus the generation of safer, more robust and more specific HbF inducers is highly desirable.

We hypothesized that a more detailed analysis of the interactions within the DRED complex might reveal new potential HbF inducers through two different possible outcomes. First, detailed information of protein-protein interactions that take place in the complexes might provide information about subunit interfaces that could provide alternative strategies for pharmacologic targeting between corepressors that have already been identified, and then this information could be used to create small molecules that would disrupt specific (possibly unique) protein interaction interfaces within the DRED complex. Second, the original mass spectrometry survey of DRED complex components (Cui et al., 2011) could easily have missed weakly or transiently interacting but nonetheless important proteins that participate in the repression mechanism, and novel γ-globin regulators might be identified that would serve as either better, or additional, HbF targets for possible therapeutic intervention. To investigate this rationale, we employed a proximity-dependent Biotin Identification strategy (BioID), a method that maps transient or low solubility protein interactions (Lambert et al., 2015) and complements our previous affinity purification strategy, to reanalyze the DRED complex in human umbilical cord blood-derived erythroid progenitor (HUDEP-2) cells, which primarily produce adult β-globin-containing RBCs *in vitro*.

Employing the BioID strategy identified the nuclear receptor corepressor-1 (NCoR1) as a TR4-interacting protein, a cofactor that was recovered at low abundance in the original affinity purification report (Cui et al., 2011). Furthermore, by employing yeast 2-hybrid assays we discovered that NCoR1 binds directly to both TR2 and TR4. We also found that NCoR1, but not TR2 or TR4, binds to enzymatic DRED corepressor subunits (e.g. DNMT1 and LSD1), suggesting that NCoR1 might serve as the scaffold upon which the TR2/TR4 DNA binding proteins indirectly convey these epigenetic modifying enzymes to act at specific chromatin sites. This hypothesis was tested by site-specific CRISPR-Cas9 mutagenesis of *NCoR1* in definitive adult HUDEP-2 cells that altered five amino acids predicted to disrupt the NCoR1 interaction interface with TR2 and TR4. The NCoR1 mutant protein as well as the DRED repressor component LSD1 failed to be recruited to their normal binding sites in the β-globin locus, confirming that NCoR1 is an adaptor for the DRED complex.

Finally, NCoR1 is known to be regulated through post-translational ubiquitination, which reduces its recruitment to specific sites in the genome (Catic et al., 2013; Mottis et al., 2013; Perissi et al., 2004; Perissi et al., 2008). Our BioID survey also identified new DRED components, including the deubiquitinase BAP1, as a potentially novel member of the complex. shRNA knockdown of BAP1 significantly reduced the recruitment of NCoR1 to sites within the globin locus, indicating that this deubiquitinase plays an important role in NCoR1 activity, and therefore in turn, DRED complex regulation. Consistent with its presumptive regulatory activity, BAP1 knockdown in HUDEP-2 cells significantly de-repressed γ-globin transcription, thereby leading to robust induction of HbF synthesis. These experiments not only detail the vital nature of previously identified protein-protein interactions that mediate γ-globin repression, but also identify novel corepressor subunits that may serve as additional therapeutic targets for future treatment of the β-globinopathies.

## Results

We previously reported that the orphan nuclear receptor heterodimer TR2:TR4 (NR2C1:NR2C2) recruited multiple epigenetic cofactors, including LSD1 (KDM1A) and DNMT1, to the promoters of the ε- and γ-globin genes, causing their transcriptional repression in adult definitive erythroid cells (Cui et al., 2011; Cui et al., 2015b; Tanabe et al., 2002; Tanabe et al., 2007). However, the precise composition as well as the number and identity of the proteins that participate in the DRED repressor complex have been only superficially defined. To address this concern, we employed BioID, a proximity-dependent labelling technique (Kim et al., 2016; Roux et al., 2012) that can detect even weak and transient protein interactions, which are often difficult to determine in standard affinity purification experiments. BioID is based on the activity of a mutant form of the prokaryotic BirA biotin ligase (BirA*). To initiate the procedure, any protein of interest is first fused to BirA*, which will, when expressed in cells, promiscuously biotinylate all proteins in its immediate *spatial* vicinity, regardless of their affinity for the protein to which the BirA* is fused. Those labeled proteins can then be collected and identified by mass spectrometry (Fig. 1A).

**Figure 1.**
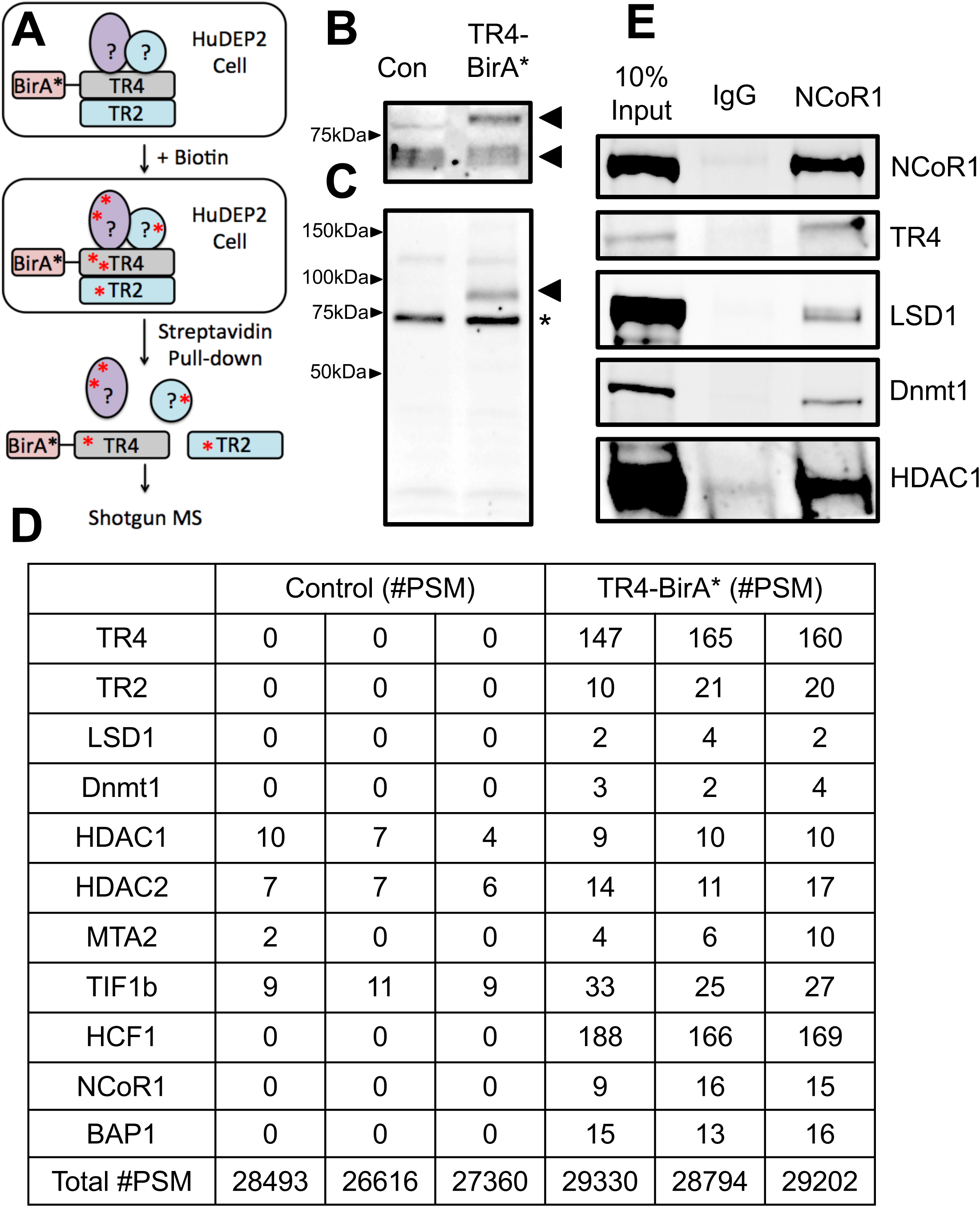
NCoR1 and BAP1 are new members of the DRED complex. (A) A schematic representation for identification of new DRED complex proteins using the BioID strategy. (B) Anti-TR4 western blotting of whole cell lysates (Cui et al. 2011. MCB). Successful generation of an active TR4-BirA* fusion protein was confirmed by demonstrating the presence of a new shifted band (upper arrow) in comparison to endogenous TR4 (lower arrow). (C) Streptavidin staining of whole cell lysates prepared from TR4-BirA*-transformed HUDEP-2 cells. The arrow indicates that the TR4-BirA* fusion protein is self-biotinylated (* indicates the presence of a background biotinylated band detected in all cell extracts). (D) Selected proteins identified in the TR4 complex (#PSM = Peptide Spectrum Matches). The total number of identified peptide sequences representing each protein is shown. (E) Co-IP confirmation that NCoR1 forms complexes with TR4, LSD1, DNMT1 and HDAC1 in HUDEP-2 cells.

To ask whether any new proteins could be identified as novel components of the DRED complex that were not co-purified in our original affinity proteomics study, BirA* was fused to the N-terminus of TR4 and then TR4-BirA* was stably transfected into HUDEP-2 cells, which upon cytokine stimulation produces mature definitive RBCs *in vitro* that primarily express adult β-globin mRNA and protein (Kurita et al., 2013). To circumvent potential artefactual effects that might result from massive TR4-BirA* forced expression, the level of fusion protein expression was maintained at close to endogenous levels (Fig. 1B; lower arrowhead = endogenous TR4, upper arrowhead = TR4-BirA*). Since the TR4 part (closest to BirA*) of the TR4-BirA* fusion protein was biotinylated by BirA* after staining with streptavidin (Fig. 1C), the data show that the TR4-BirA* fusion protein, when expressed at roughly endogenous levels, is able to identify other proteins in its immediate vicinity.

After incubation in the presence of biotin, the TR4-BirA* as well as parental HUDEP-2 cells were lysed, and the biotinylated proteins were isolated and identified by mass spectrometry. Among the output of total peptide spectrum matches (#PSM) from three independent experiments, all of the DRED complex proteins, including LSD1 and DNMT1, that were identified previously by affinity purification (Cui et al., 2011) were confirmed as TR4 (either direct or indirect) interacting proteins. In addition to confirming the identity of proteins that were previously identified in the complex, the analysis also revealed new proteins that were in close proximity to the TR4 fusion protein (Fig. 1D and Supplemental Table 1). Subsequent studies described below demonstrated that two of the BioID-labeled proteins, NCoR1 and BAP1, play critical roles in γ-globin repression.

### NCoR1 is a key adaptor protein in the DRED complex

The nuclear receptor corepressor-1 (NCoR1) has been shown to recruit HDACs into large macromolecular complexes with thyroid hormone receptor, retinoic acid receptor as well as other non-nuclear receptor transcription factors to mediate transcriptional repression of target genes (Mottis et al., 2013; Perissi et al., 2008). In this regard, we hypothesized that NCoR1 might serve as the adaptor between TR2/TR4 and other repressive components of DRED complex. To test this hypothesis, protein complexes in wild-type HUDEP-2 nuclear extracts were first immune precipitated using an anti-NCoR1 antibody followed by western blotting using antibodies that recognize LSD1, DNMT1, HDAC1 or TR4 (Fig. 1E). These co-IP data indicate that endogenous NCoR1 interacts with endogenous TR4 as well as each of the other epigenetic modifying enzymes in HUDEP-2 cells, consistent with the hypothesis that NCoR1 might serve as the scaffold upon which the DRED repressor complex is assembled.

To address whether NCoR1 might act as the scaffold for TR2 and TR4 to recruit corepressor enzymes, we employed yeast two hybrid (Y2H) assays to examine the interactions between these proteins using fragments of NCoR1 as bait. Successful expression of LexA-bait and VP16-prey fusion proteins was confirmed by western blots of yeast protein extracts probed with anti-LexA or anti-VP16 antibodies (Supplemental Fig. 1). Since NCoR1 is encoded by 2453 amino acids, we generated three separate, overlapping fragments of NCoR1 to use as bait proteins (Table 1). Those three fragments contain, respectively: repressive domain 1 (RD1); repressive domains 2 plus 3 (RD2/3); or the carboxy-terminal (C) domain (Mottis et al., 2013).

**Table 1.** Yeast 2 hybrid (Y2H) studies of NCoR1 binding to other candidate proteins within the DRED complex. The relative scores for protein-protein interactions between fragments of NCoR1 and previously identified members of the DRED complex based on the induction of the HIS3 reporter gene in Y2H assays that allow L40 yeast to grow in media lacking histidine. The scores indicate interactions from no growth (-) to robust growth (5+). * = N-terminal amino acids 2-757; ** = Mi2β amino acids 667-1295; # = C-terminal amino acids 1630-2454; ## = C-terminal amino acids 2004-2468.

| Prey<br>(VP16) | Bait (LexA) |  |  |  |  |  |
| --- | --- | --- | --- | --- | --- | --- |
|  | NCOR1 |  |  | TR2 | TR4 | Sin3A |
|  | RD1 | RD 2/3 | C |  |  |  |
| VP16 | - | - | - | - | - | - |
| TR2 | - | - | 3+ | 5+ | 5+ | 3+ |
| TR4 | - | - | 4+ | 5+ | 5+ | 3+ |
| LSD1 | 2+ | - | 2+ | - | - | - |
| DNMT1* | 1+ | - | - | - | - | - |
| DNMT1 | - | - | - | - | - | - |
| Rcor1 | 1+ | - | - | - | - | - |
| Rcor2 | - | - | - | - | - | - |
| HDAC1 | - | - | - | - | - | - |
| HDAC3 | - | - | 2+ | - | - | - |
| TRIM28 | - | - | - | - | - | - |
| CtBP1 | - | - | 2+ | - | - | - |
| Sin3A | - | - | 5+ | 1+ | 2+ | - |
| Mi2$\beta^{**}$<br>(CHD4) | - | - | <1+ | - | - | - |
| NCOR1# | - | - | 5+ | 1+ | 2+ | 4+ |
| SMRT1## | - | - | - | - | - | - |

When fragments of NCoR1 were tested in Y2H experiments, the RD1 domain of NCoR1 was found to interact with LSD1, the N-terminus of DNMT1 and Rcor1 (CoREST; Table 1). Furthermore, the C-terminal domain of NCoR1 was also found to interact with several corepressors including LSD1, HDAC3, CtBP1, Sin3A and mi2β (one of multiple proteins that constitute the mammalian NuRD complex) as well as with TR2 and TR4. None of the corepressors that we tested interacted with the NCoR1 RD2/3 domain (Table 1).

Interestingly, among all of the tested corepressor proteins, only NCoR1 (C-terminal) and Sin3A bound directly to TR2/TR4 (when TR2 or TR4 were used as Y2H bait; Table 1), indicating that NCoR1 (and/or Sin3A) serves as an adaptor platform between TR2/TR4 and the other DRED complex subunits. Based on these data, we then addressed the possibility that Sin3A might be an additional or alternative DRED scaffold. However, when Sin3A was used as bait in other Y2H experiments, we found that TR2, TR4 and NCoR1, but none of the epigenetic modifying corepressor enzymes, bound to Sin3A (Table 1), suggesting either that Sin3A is not an adaptor in the DRED complex or that it mediates interactions with additional partner proteins that have not yet been identified. Notably, the C-terminal domain of SMRT (silencing mediator of retinoic acid and thyroid hormone), which contains nuclear receptor interaction domains and also can serve in scaffold interactions between nuclear receptors and epigenetic co-repressors (Mottis et al., 2013), does not directly interact with either TR2 or TR4 (Table 1), indicating that NCoR1 fulfils a unique requirement as the adaptor between TR2/TR4 and the DRED co-repressor enzymes. Taken together, the data were consistent with a hypothesis regarding the protein constituents that might be minimally required to generate the large DRED repressor: NCoR1 serves as the central adaptor protein in which different domains recruit both the DNA binding components (TR2 and TR4) as well as critical epigenetic modifying enzymes into the repressor complex.

### TR4 and NCoR1 genomic binding site occupancy overlaps in K562 cells

The data reported indicates that NCoR1 might serve as the scaffold between TR2/TR4 and other DRED complex proteins. If this hypothesis is correct, then NCoR1 and TR2/TR4 ChIP-seq peaks would be predicted to significantly overlap in the erythroid genome. To address this hypothesis, we examined the ChIP-seq signatures of NCoR1 (−1 and −2), EGFP-TR2 and EGFP-TR4 in K562 cells (a human embryonic myeloerythroid cell line) from the ENCODE database (http://genome.ucsc.edu/encode/downloads.html). The common peaks shared between two different datasets (NCoR1-1, NCoR1-2) were accumulated as NCoR1 peaks to calculate NCoR1 and TR2/TR4 peak overlap. Among the total of 17,417 NCoR1 peaks, 6195 (35.5%) and 6932 (39.8%) co-localized with TR2 and TR4 respectively, and 4538 (26%) peaks overlapped both TR2 and TR4 (Fig. 2A). Additionally, among the total of 22,243 TR2 and 26,557 TR4 peaks, 6195 (27.9%) and 6932 (26.1%) were found to be co-occupied by NCoR1 (Fig. 2A).

**Figure 2.**
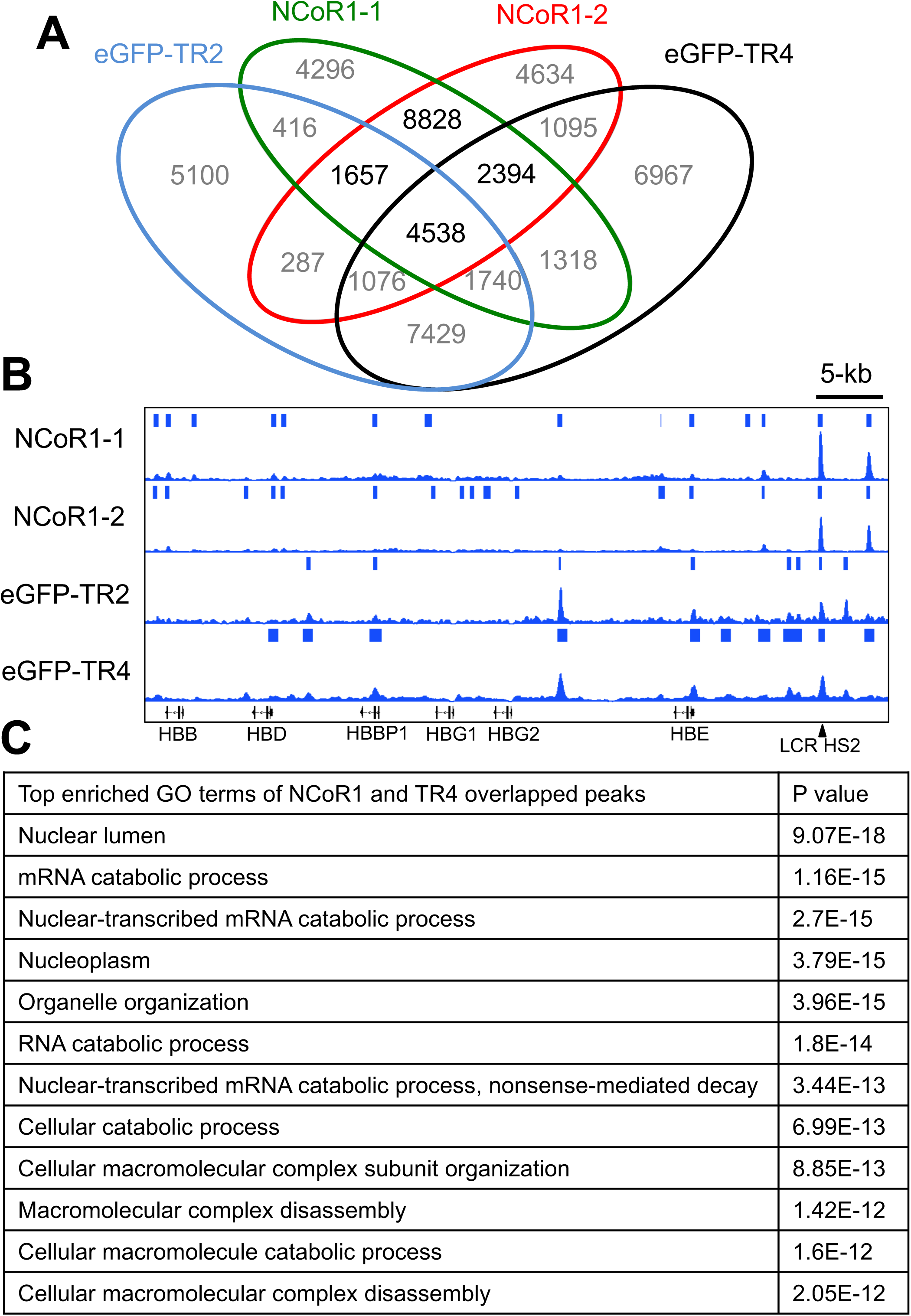
TR4 and NCoR1 significantly overlap in genome wide binding site distribution. (A) Venn diagram of TR4, TR2 and NCoR1 ChIP-seq peaks in human erythroleukemia K562 cells. (B) Representative NCoR1, TR2 and TR4 overlapping peaks in the β-globin locus. (C) Top enriched GO terms extracted from the TR4, NCoR1-1 and NCoR1-2 overlapping peak-associated genes.

The β-globin locus and the genes that regulate RNA catabolism and cell cycle were chosen as representative loci for association with all of these proteins, and there are clearly overlapping as well as unique binding site signatures (Fig. 2B and Supplemental Fig. 2). The strongest overlap among all three Chipped proteins in the β-globin locus in K562 cells was at a DR2 element (GC<u>TGACCAC</u>C<u>TGACTA</u>AA) in LCR HS2. Taken together, these ChIP-seq data indicate that NCoR1 and TR2/TR4 extensively overlap in the erythroid genome, consistent with the concept that these proteins often function together in the DRED complex for a significant fraction of time (e.g. LCR HS2; Fig. 2B). Among the top GO enrichment pathway terms among the peaks shared by NCoR1 and TR4 (Fig. 2C), there are multiple terms describing enrichment of proteins involved in macromolecular catabolism and cell organelle organization, suggesting that the DRED complex likely plays additional, unexplored roles in these pathways.

### NCoR1-ID3 binds to the ligand-binding domains of TR2 and TR4

To begin to detail the potential mechanisms of transcriptional repression elicited by the DRED complex, we characterized specific protein-protein interaction interfaces between TR2, TR4 and NCoR1. A series of TR2 or TR4 truncations was generated and tested by Y2H in order to map the domains that mediate their interactions with the C-terminal domain of NCoR1 (NCoR1-C; Table 1). The expression of LexA-bait and VP16-prey fusion proteins was confirmed by western blots using anti-LexA and anti-VP16 antibodies (Supplemental Fig. 1). These results show that the ligand binding domains (LBD; EF domains; Table 2) of both TR2 and TR4 were responsible for the interaction with NCoR1-C in a manner similar to other nuclear receptor/NCoR1 associations (Jepsen and Rosenfeld, 2002), and this interaction does not require the N-terminal domain (regions A/B), DNA-binding domain (region C) or the hinge region (region D) of either orphan receptor (Table 2). Additionally, the N-terminal domain (A/B) of TR4, but not TR2, is capable of independently interacting with NCoR1-C; thus two different domains of TR4 can interact with NCoR1-C.

**Table 2.** Identification of the binding interface between NCoR1 and TR2 or TR4. Fragments of TR2 or TR4 were used as baits in potential Y2H interactions with NCoR1 wild-type and mutant interaction domains (ID3*) based on the induction of a HIS3 reporter gene that allows L40 yeast to grow in medium lacking histidine. The scores range from no growth (-) to robust growth (5+). ND; not determined. FL = full length; ND = not determined; * = IDVII to ADAIA mutation

|  | TR2 |  |  |  |  | TR4 |  |  |  |  |
| --- | --- | --- | --- | --- | --- | --- | --- | --- | --- | --- |
|  | FL | A/B | C | D | E/F | FL | A/B | C | D | E/F |
| NCoR1-C | 3+ | - | - | - | 4+ | 4+ | 3+ | - | - | 5+ |
| NCoR1-ID1 | - | ND | ND | ND | ND | - | - | ND | ND | ND |
| NCoR1-ID2 | - | ND | ND | ND | ND | - | - | ND | ND | ND |
| NCoR1-ID3 | 1+ | - | - | - | 3+ | 3+ | 3+ | - | - | 5+ |
| NCoR1-ID3* | ND | ND | ND | ND | - | - | <1+ | ND | ND | - |

The C-terminus of NCoR1 has three interaction domains (ID1-ID3; Table 1) that were reportedly responsible for its interaction with other nuclear receptors (Cohen et al., 2001). Each of the IDs contains a corepressor-nuclear receptor (CoRNR) box bearing the 5 amino acid consensus sequence (I/L)XX(V/I)I (where X is any amino acid) that preferentially interacts with nuclear receptors (Hu and Lazar, 1999; Perissi et al., 1999). To assess which ID(s) of NCoR1 interact with the LBDs of TR2 and TR4, three truncated fragments containing each of the NCoR1-IDs were tested in Y2H assays. As shown in Table 2, the LBDs of TR2 and TR4 specifically interacted with NCoR1-ID3, demonstrating an ID3-preference for TR2/TR4:NCoR1 interactions. Furthermore, mutation of three amino acids in the ID3-CoRNR box (IDVII to ADAIA) fully abolished interactions with both TR2 and TR4 (Table 2), demonstrating a specific requirement for the ID3-CoRNR box for interactions between the DRED orphan nuclear receptors and NCoR1-ID3. Taken together, these results clearly demonstrate that NCoR1 uses a CoRNR box in its ID3 domain to specifically bind to sequences in the LBDs of both TR2 and TR4.

### NCoR1 is the adaptor for the DRED complex in (human adult definitive) HUDEP-2 cells

By identifying the amino acids in the NCoR1 ID3 domain that are critical for TR2/TR4 interaction, we next employed CRISPR-Cas9 editing to generate site directed mutants of NCoR1 ID3 in HUDEP-2 cells that were predicted to block interactions with wild type TR2/TR4. Two NCoR1 AAAAA mutant clones were established and the site-specific mutagenesis of both clones was verified by Sanger sequencing (Fig. 3A). Consistent with the Y2H data shown in Table 2, the mutation of the five amino acids that comprise the CoRNR box completely disrupted NCoR1 interaction with TR4 in HUDEP-2 cells, whereas the interaction between the mutant NCoR1 protein and LSD1 is unperturbed (Fig. 3B).

**Figure 3.**
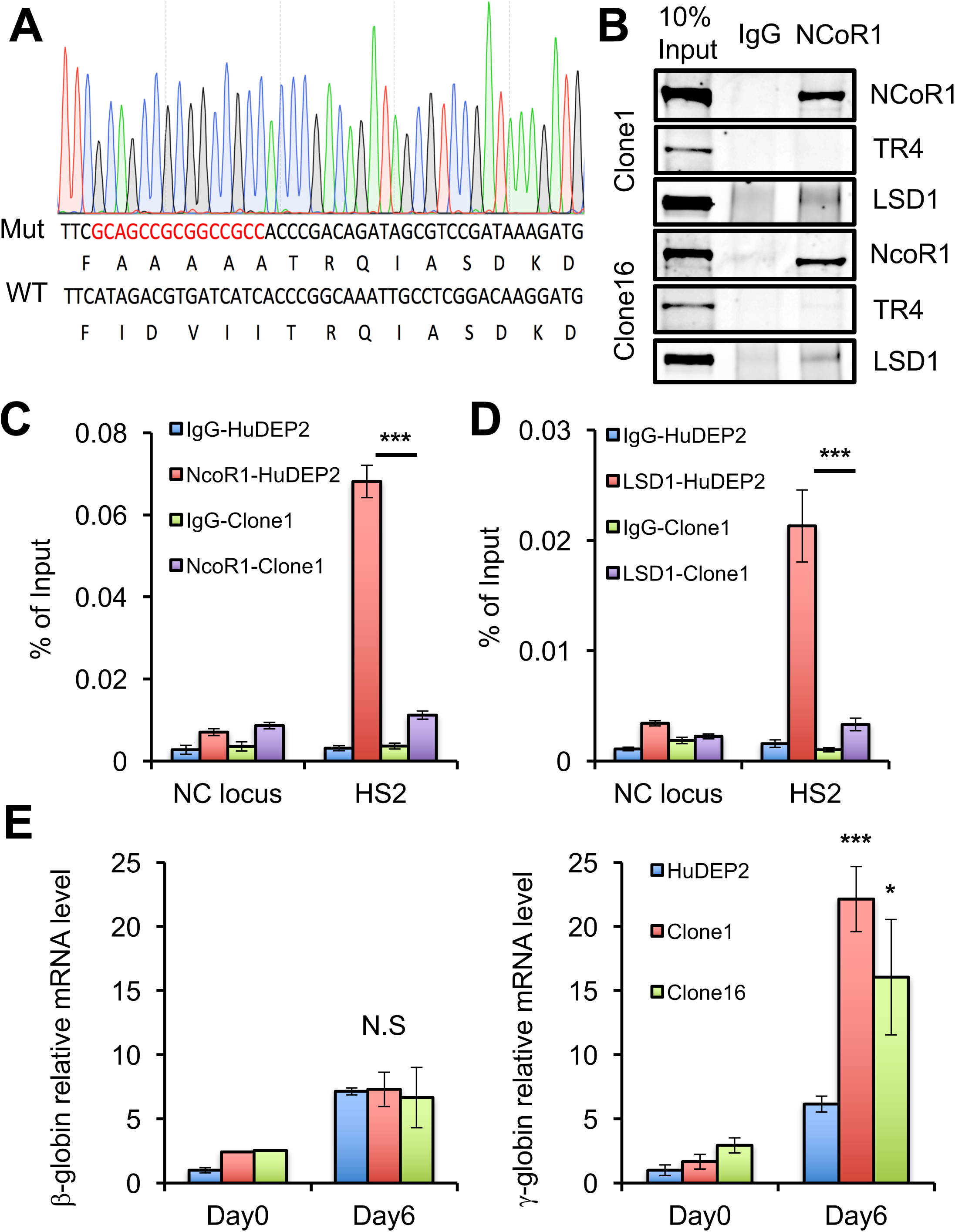
NCoR1 is the direct adaptor between TR2/TR4 and other DRED complex corepressors. (A) Design and generation of an NCoR1 mutant with deficient TR4 interaction. Red letters represent the mutant nucleic acid sequence generated by CRISPR targeting that destroys the ID3-CoRNR box (Hu and Lazar, 1999; Perissi et al., 1999). (B) Confirmation that the TR4:NCoR1 interaction is abolished whereas the LSD1:NCoR1 interaction is maintained in two individual NCoR1 CoRNR mutant clones by co-IP in HUDEP-2 cells. (C) A CRISPR/Cas9 generated NCoR1 mutant HUDEP-2 cell line fails to be recruited to an endogenous TR2/4 binding site in the β-globin locus. (D) LSD1 occupancy is significantly reduced at the HS2 TR2/4 binding site in the β-globin locus in the absence of a robust NCoR1:TR4 interaction in undifferentiated HUDEP-2 cells. Blue bar, WT HUDEP-2 with IgG control; Red bar, WT HUDEP-2 cells with anti-NCoR1 or anti-LSD1; Green bar, NCoR1 mutant HUDEP-2 with IgG control; Purple bar, NCoR1 mutant HUDEP-2 cells with anti-NCoR1 or anti-LSD1. (E) Disruption of the TR2/TR4:NCoR1 interaction within the DRED complex derepresses γ-globin transcription in HUDEP-2 cells. (Left) β-globin and (Right) γ-globin relative mRNA abundances in undifferentiated or differentiated (for 6 days) wild-type or NCoR1-mutant HUDEP-2 cells. Data are shown as the mean ± SD from three independent experiments. (\**p*<0.05; \*\*\**p*<0.001; unpaired Student’s *t*-test).

In the β-globin locus, wild-type NCoR1 binds prominently to HS2 in the LCR (Fig.s 2B and 3C). However, recruitment of the AAAAA mutated NCoR1 to its most prominent β-globin locus binding site in HS2 was significantly reduced (Fig. 3C), indicating that TR2/4 plays a central role in NCoR1 recruitment to globin locus chromatin. Consistent with the concept that NCoR1 is the adaptor between TR2/4 and multiple corepressor enzymes, LSD1 recruitment at HS2 of the locus was also significantly reduced in NCoR1 mutant HUDEP-2 cells (Fig. 3D). Taken together, these data confirmed that NCoR1 serves as a primary adaptor to combine TR2/4 with DRED corepressor proteins.

### Disruption of the TR2/4:NCoR1 interaction derepresses γ-globin transcription

The DRED complex is critical for γ-globin repression in adult red blood cells (Suzuki et al., 2014) and NCoR1 knockdown by shRNA in CD34+ cell erythroid differentiation cultures has been shown to induce γ-globin mRNA synthesis (Xu et al., 2013). To test for possible functional deficiencies that are due to NCoR1 loss of function in γ-globin repression, we determined globin mRNA levels in undifferentiated (day-0) or differentiated (day-6) HUDEP-2 cells bearing NCoR1 mutations that disrupt its interaction with TR2/4 (Fig. 3E). Using either independent NCoR1 mutant HUDEP-2 clone, both β-globin and γ-globin expression were very similar in wild type and mutant cells in the absence of differentiation induction. However, after 6 days of erythroid differentiation induction, while β-globin expression was essentially unchanged from its level in wild type cells, γ-globin mRNA in the mutant clones increased by 2- to 3-fold, demonstrating that disruption of the TR2/TR4:NCoR1 interface specifically de-represses γ-globin expression in differentiated erythroid progenitor cells.

### Loss of the NCoR1 ID3 domain induces embryonic murine globin transcription

Among the three interaction domains of NCoR1 that specifically bind to various nuclear receptors, we showed that only ID3 is critical for NCoR1 interaction with both TR2 and TR4 (Table 2). To further assess the functional significance of TR2/4 binding of NCoR1 on γ-globin repression *in vivo*, we took advantage of a conditional NCoR1 mouse mutant that bears loxP sites flanking exons 38-41 (encoding the ID3 and ID2 domains; Supplemental Fig. 3)(Astapova et al., 2008). Upon Cre-mediated recombination, the NCoR1 gene is predicted to encode a protein missing ID2 and ID3. The NCoR1 mutant mice were bred to Mx1-Cre mice (Kuhn et al., 1995) to induce deletion of NCoR1 ID2/3 in all hematopoietic lineages. Since the murine *εy* and *βh1* genes bear DR sequences at the same relative positions in their promoters as the human ε- and γ-globin genes, respectively, induction of *εy* and *βh1* have become reliable reflections of the regulatory activity of their human counterparts.

The effect of deletion of NCoR1 ID2/3 was evaluated in erythroid progenitor cells isolated from E14.5 fetal of livers NCoR1^flox/flox^:Tg^Mx1Cre^ and NCoR1^flox/flox^ (control) embryos (Supplemental Fig. 3A). Interferon-α (IFN-α) was added to both progenitor cell cultures to activate expression of the Cre transgene. After induction, 70% of the mutant *ncor1* alleles were found to be deleted in the erythroid cells of IFN-α-treated *ncor1* mutant mice, whereas no deletion was detected in control animals (Supplemental Fig. 3B). The 70% recombination of *ncor1* locus led to a 2- to 3-fold induction of murine *βh1* (the γ-globin homolog) transcription (Supplemental Fig. 3C) after induction of erythroid differentiation, showing that NCoR1 ID2/3 contributes to embryonic *βh1* gene repression in mice. The data show that ID2/3-deleted NCoR1 leads to induced embryonic globin mRNA *ex vivo* in erythroid cells.

### BAP1 regulates β-globin locus NCoR1 recruitment

In addition to the previously defined DRED complex corepressors and NCoR1 that were identified here using BioID, we detected strong HCF1 association with TR4 (Fig. 1D), which was in agreement with the interactome of TR2/TR4 that was previously demonstrated in murine MEL cells (Cui et al., 2011). Interestingly, the well-characterized HCF1-interacting tumor suppressor BRCA1 associated protein-1 (BAP1) was also identified as a new component in the DRED complex (Fig. 1D). BAP1 is a nuclear-localized ubiquitin carboxyl-terminal hydrolase (Dey et al., 2012; Lee et al., 2014; Qin et al., 2015; Zarrizi et al., 2014), which has been shown to stabilize nuclear proteins through its deubiquitinase activity. Among the multiple corepressors already identified in the DRED complex, NCoR1 has been explicitly shown to be regulated by ubiquitination (Catic et al., 2013; Mottis et al., 2013; Perissi et al., 2004; Perissi et al., 2008), and the ubiquitination of NCoR1 has been proposed to modulate its stability and genome recruitment through proteasome degradation (Catic et al., 2013). Based on previous studies as well as our demonstration that both NCoR1 and BAP1 interact with TR4 (Fig. 1D), we investigated the possibility that BAP1 might regulate NCoR1 activity through its deubiquitinase activity.

To test this hypothesis, we immune precipitated HCF1 from wild-type HUDEP2 nuclear extracts and probed those immune complexes by western blotting with TR4, NCoR1 and BAP1 antibodies. The co-IP experiments indicated that all four of these proteins can be found in complex (Fig. 4A), consistent with the proteomics data. Next, the HCF1 co-IP was performed using the AAAAA NCoR1 mutant HUDEP-2 cell clone nuclear extracts in which the TR4 interaction with NCoR1 has been disrupted (Table 2 and Fig. 3B). In these cells, mutant NCoR1 is still able to bind HCF1 (Supplemental Fig. 4), indicating that NCoR1:HCF1 binding is not dependent on TR4:NCoR1 interaction.

**Figure 4.**
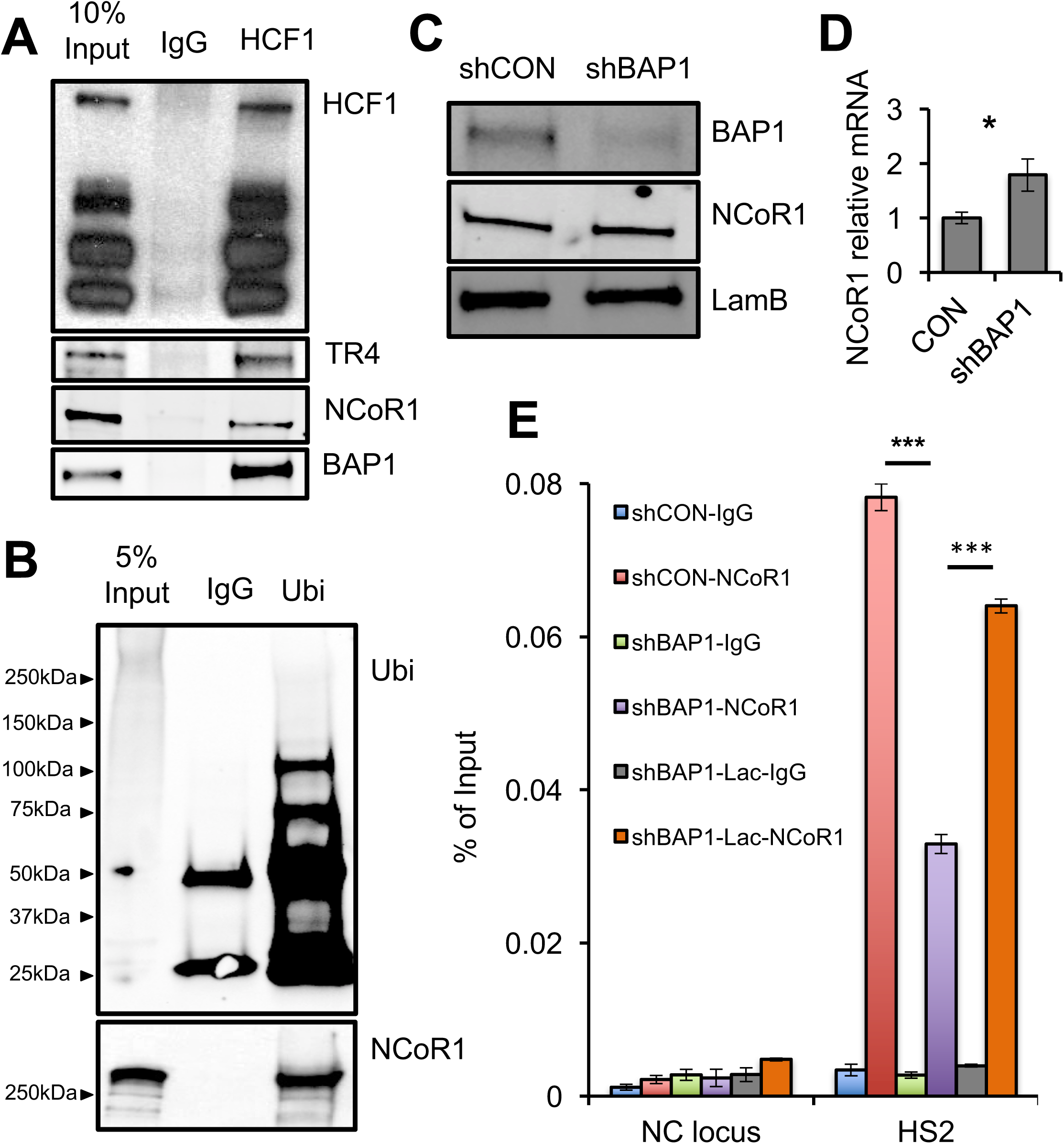
HCF1 and BAP1 are new components of the DRED complex and regulate NCoR1 activity in the β-globin locus. (A) Immune precipitated HCF1 was interrogated on western blots using antibodies recognizing HCF-1, TR4, NCoR1 or BAP-1 in wild type HUDEP-2 cells. (B) Immune precipitated anti-ubiquitin extracts from wild type HUDEP-2 cells were interrogated with anti-ubiquitin (top) or NCoR1 (bottom panel) antibodies. (C) shRNA knockdown of BAP-1 in HUDEP-2 cells only marginally affects NCoR1 or lamB protein abundance. (D) NCoR1 mRNA level increased by only 1.7-fold after BAP1 knockdown. (E) NCoR1 occupancy at the TR2/4 binding site in HS2 is significantly reduced after BAP-1 knockdown in undifferentiated HUDEP-2 cells, and the reduced NCoR1 occupancy could be partially rescued by treatment with the proteasome inhibitor, lactacystin. Blue bar, control scrambled shRNA infected HUDEP2 with IgG control; Red bar, control HUDEP2 cells with anti-NCoR1; Green bar, BAP1 knockdown HUDEP2 with IgG control; Purple bar, BAP1 knockdown HUDEP2 cells with anti-NCoR1; Gray bar, BAP1 knockdown HUDEP-2 after 3 hours of treatment with 25 µM lactacystin, IgG control; Orange bar, NCoR1 detection in BAP1 knockdown HUDEP-2 cells after 3 hours of treatment with 25 µM lactacystin. Data are shown as the means ± SD from three independent experiments (\**p*<0.05; \*\*\**p*<0.001; unpaired Student’s *t*-test).

To examine whether NCoR1 becomes ubiquitinated in wild type HUDEP-2 cells, we next immune precipitated nuclear extracts using an anti-ubiquitin antibody followed by western blotting with anti-NCoR1. When compared with control IgG immune precipitates, the anti-ubiquitin antibody co-precipitates a significant amount of NCoR1, indicating that some fraction of the nuclear NCoR1 is ubiquitinated in HUDEP-2 cells. The data suggests that the ubiquitination status of NCoR1 might be important for its function (Fig. 4B).

To test whether BAP1 and ubiquitination alter NCoR1 recruitment to chromatin [as previously reported (Catic et al., 2013)], we reduced the abundance of BAP1 mRNA using an shRNA-containing lentivirus followed by NCoR1 ChIP assays at the β-globin locus. The efficiency of BAP1 knockdown was confirmed by western blotting (Fig. 4C). We found that reduced BAP1 abundance does not significantly alter the level of NCoR1 protein (Fig. 4C), whereas the NCoR1 mRNA level slightly increased (Fig. 4D). When compared to infection with a control scrambled shRNA, BAP1 knockdown significantly reduced the recruitment of NCoR1 at its most prominent binding site in the globin locus at LCR HS2 (Fig. 4E, purple vs red bars).

Ubiquitination of NCoR1 destabilized its occupancy on chromatin, presumably by acting through proteasome-mediated protein degradation. Treatment with the proteasome inhibitor, Lactacystin (Lac), has been reported to significantly enhance NCoR1 recruitment at specific chromosomal loci by preventing its degradation (Catic et al., 2013). To test whether BAP1 knockdown would reduce NCoR1 recruitment to HS2 through a proteasome-mediated process, we first treated uninfected HUDEP-2 cells with Lac for 3 hours, which significantly enhanced accumulation of ubiquitinated protein in a dose-dependent manner, while total cellular NCoR1 abundance was unchanged (Supplemental Fig. 5). When anti-BAP1 shRNA lentivirus-infected HUDEP-2 cells were treated with Lac for 3 hours, NCoR1 occupancy at HS2 was rescued (Fig. 4E, orange vs purple bars), indicating that proteasome-mediated NCoR1 degradation plays a central role in its chromatin occupancy. Taken together, these data suggest that both BAP1 and NCoR1 are involved in DRED complex activity and that BAP1 is a key regulator of NCoR1 recruitment to specific chromatin sites in the β-globin locus.

### Reduced BAP1 expression robustly induces γ-globin transcription

Since BAP1 was identified here as a novel regulator of DRED complex activity that appears to be important for NCoR1 recruitment to β-globin locus HS2 (Fig. 4E), we next asked whether BAP1 knockdown would functionally affect fetal globin gene expression. To do so, mRNA abundances were assessed in undifferentiated (day-0) or differentiated (day-6, same as in Fig. 3E) BAP1 shRNA- and control shRNA-infected HUDEP-2 cells.

BAP1 gene expression increased during erythroid differentiation (day 6 vs day 0) in control (scrambled) shRNA-infected HUDEP-2 cells, whereas BAP1 mRNA was reduced to about 30% of controls in shBAP1-infected cells (Fig. 5A, left panel). At day-0, BAP1 knockdown induced β-globin mRNA levels by 4-fold and γ-globin by 43-fold (Fig. 5A, middle and right panels). After 6 days of erythroid differentiation, BAP1 knockdown further induced β-globin expression slightly, whereas γ-globin transcription was massively induced (approximately 90-fold; Fig. 5A, right panel). The significant de-repression of γ-globin synthesis was confirmed by HPLC monitoring of hemoglobin production in the BAP1 knockdown cells (Figw. 5B); the percentage of HbF increased from 0.1% (shCON) to 13.5% (shBAP1) after 10-days of erythroid differentiation induction. Considering that the shRNA knockdown efficiency of BAP1 was around 70% (Fig. 4C), the data indicate that this newly discovered DRED complex regulator, BAP1, plays a vital role in fetal globin gene repression, and that inhibition of BAP1 deubiquitinase activity may serve in the future as a novel therapeutic target for γ-globin induction.

**Figure 5.**
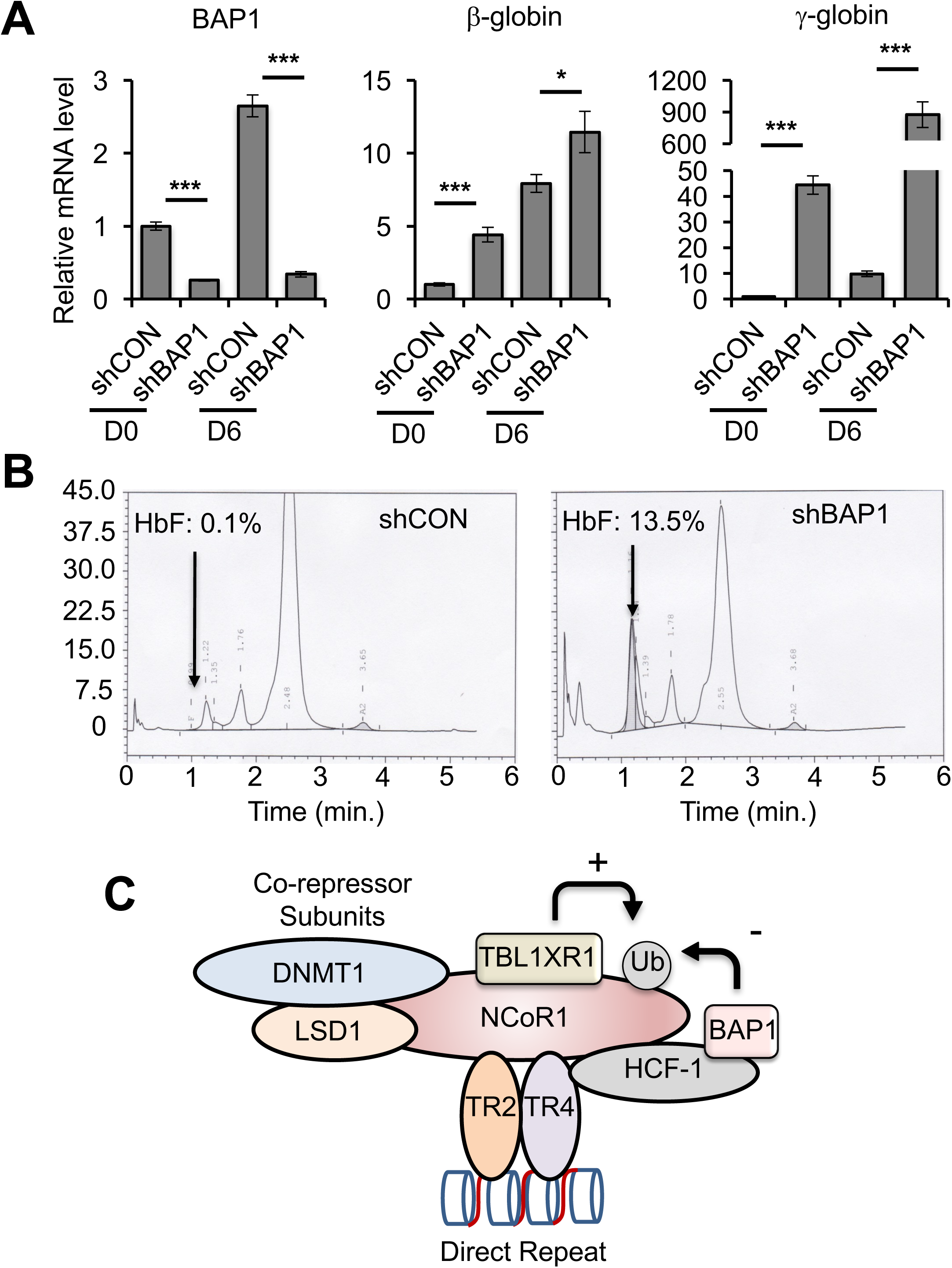
Reduced BAP1 expression robustly induces γ-globin transcription. (A) The relative mRNA abundance of BAP1, β-globin and γ-globin in undifferentiated (D0) or differentiated (D6) HUDEP-2 cells. (B) Reduced BAP1 expression significantly induces HbF after 10 days of HUDEP-2 cells differentiation induction. (C) Schematic summarizing the observations that NCoR1 is the scaffold on which TR2/TR4 and the corepressor epigenetic enzymatic subunits of the DRED complex are assembled. We hypothesize that the ubiquitination of NCoR1 by TBL1XR1, or its deubiquitination by BAP1, regulates a dynamic homeostasis of NCoR1 recruitment or retention in the DRED complex and subsequent globin gene repression and de-repression (\**p*<0.05; \*\*\**p*<0.001; unpaired Student’s *t*-test).

## Discussion

In this study, we found that NCoR1 serves as a central scaffold upon which the active DRED repressor is assembled. We also showed that a novel DRED repressor subunit, deubiquitinase BAP1, regulates site-specific NCoR1 recruitment within the β-globin locus and that BAP1 reduction leads to enormously increased γ-globin mRNA and protein induction (Fig. 5C).

Several lines of evidence demonstrate that BAP1 regulates NCoR1 recruitment through its deubiquitinase activity. First, NCoR1 is ubiquitinated in HUDEP-2 cells (Fig. 4B) and the ubiquitination of NCoR1 followed by proteasome degradation was previously reported to be vital for the regulation of NCoR1 transcriptional activity (Catic et al., 2013). Second, BAP1 is found in a complex with NCoR1 (as well as with TR4 and HCF1; Fig. 4A) and knockdown of BAP1 mRNA reduced NCoR1 recruitment at β-globin locus-associated sites and could be rescued by proteasome inhibitor treatment (Fig. 4E). Third, although BAP1 erythroid ChIP-seq information is not yet available, HCF1 [to which the majority of cellular BAP1 appears to be bound (Fig. 4A), and is found in complex with a majority (85%) of BAP1 in bone marrow-derived macrophage (Dey et al., 2012)] co-localized with NCoR1 genome wide as well as in the globin locus in K562 erythroleukemia cells (from ENCODE ChIP-seq studies). Additionally, the F-box-like/WD repeat-containing protein TBL1XR1 that is responsible for NCoR1 ubiquitination (Mottis et al., 2013; Perissi et al., 2004; Perissi et al., 2008) also co-localized with NCoR1 and HCF1 as well as with TR2/TR4 at sites in the β-globin locus in K562 cells (ChIP-seq in ENCODE). Taken together, the available data suggest that in vivo homeostasis of NCoR1 is achieved by an opposing ubiquitination/deubiquitination balance, and that this likely plays a key regulatory role in NCoR1 activity in erythroid cells (Fig. 5C).

mRNA stability is an important mechanism to regulate protein abundance. Globin mRNA stability is a critical determinant of protein abundance in normal erythropoiesis since the long half-life of these mRNAs is fundamental for the continuous translation of globin proteins, which is critical during later stages of erythropoiesis when transcription becomes arrested and nuclei are shed (Waggoner and Liebhaber, 2003). In contrast, destabilization of globin mRNA by naturally occurring mutations such as anti-termination signals [TAA to CAA in the α2-globin gene leading to accelerated mRNA decay (Morales et al., 1997)] results in α-thalassemia. Under physiological conditions, it is possible that nonsense transcripts could be generated and that the cell is able to destabilize these aberrant transcripts through various pathways including nonsense-mediated decay (Palacios, 2013), indicating that mRNA catabolism could play a role in erythropoiesis. In this regard, it is interesting to note that the GO terms shown in Figure 2 (and representatives in Supplemental Figure 2) suggest that the genomic sites of overlap between TR4 and NCoR1 are enriched in multiple pathways associated with RNA catabolism and nonsense-mediated mRNA decay, suggesting that TR4 and NCoR1 might affect erythropoiesis through transcriptional regulation of pathways that control mRNA stability.

In a previous study, we found that the TR4 promotes erythroid cell proliferation in mouse bone marrow-derived erythroid progenitors (Lee et al., 2017). In this regard, it is interesting to note that the both of the HUDEP-2 clones harboring an NCoR1 mutation that leads to failure to interact with TR2/TR4 grow significantly slower than parental cells (data not shown), suggesting that NCoR1 could mediate cell cycle regulatory activity of TR4 as the DRED complex adaptor. By analyzing the TR2/TR4 and NCoR1 ChIP-seq database in K562 cells we found co-occupancy of all three proteins near the promoters of E2F family transcription factors, critical regulators of G1 to S phase transition during the cell cycle (Supplemental Fig. 2). This suggests that TR4 might regulate cell proliferation through direct recruitment of NCoR1 and subsequent transcriptional control of E2F family members.

The switch from corepressor to co-activator function is an important gene regulatory mechanism for nuclear receptors (Perissi et al., 2010). Based on this fact for which many examples exist and the model proposed in this study, there are intriguing questions that remain. First, what protein(s) (if any) constitute the co-activator form of TR2/TR4 after depleting cells of NCoR1? A previous study suggested that PGC-1α/β might be such co-activator candidates (Cui et al., 2014). PGC-1α serves as the co-activator for several other nuclear receptors such as PPARγ (Lin, 2009), and PGC-1α/β are in fact recruited in the mouse β-globin locus, where TR2/TR4 bind and the interaction between TR2/TR4 and PGC-1α/β was confirmed by co-immunoprecipitation in erythroid cells (Cui et al., 2014). Furthermore, compound mutants in the PGC-1α/β genes impaired murine β-type globin gene expression, which is induced by disrupting TR2/TR4 and NCoR1 interaction (Fig. 3E), suggest that NCoR1 and PGC-1α/β share a mutually antagonistic relationship. Taken together, these data suggest that PGC-1α/β might serve as candidate coactivators that replace ubiquitin-mediated turnover of NCoR1 to generate a TR2/4 activator from the DRED repressor form.

An additional question regards the mechanism that could allow or promote the exchange of NCoR1 for coactivators. A study showing that ubiquitination of NCoR1 by TBL1XR1 favors the switch from corepressors to coactivators has been reported, which may be the underlying mechanism responsible for a cofactor switch (Mottis et al., 2013). NCoR1 has been shown to bind to unliganded nuclear receptors, and this interaction is lost upon ligand binding (Perissi et al., 2010). In this regard, it is interesting to note that non-physiological concentrations of vitamin A (or more likely a related molecule) has been promoted as a potential TR4 ligand from in vitro analysis (Zhou et al., 2011). If there is a high affinity ligand for either or both TR2 and TR4, such a fact could have significant clinical potential, since ligand antagonists might be identified that could induce γ-globin transcription toward the aim of benefiting patients with β-globinopathies.

In this study, we discovered that a loss of NCoR1 recruitment to β-globin locus sites induced fetal globin gene expression by 2- to 3-fold, whereas a 70% reduction in BAP1 enzyme abundance induces γ-globin synthesis 90-fold, indicating that there are other potential γ-globin inductive mechanisms that are regulated by BAP1. Of note, one other potent γ-globin repressor, bcl11a, has also been shown to be complexed with NCoR1 (Xu et al., 2013), suggesting the intriguing possibility that BAP1 activity regulates multiple γ-globin repressors. To delve more deeply into the underlying mechanism by which BAP1 mediates repression of γ-globin synthesis, it would be of great significance to determine the identity of additional BAP1-target proteins in erythroid cells. Detailed investigation of this issue is of potential clinical significance since increased γ-globin synthesis is known to mitigate the symptoms of the β-globinopathies (Ngo et al., 2012; Noguchi et al., 1988; Weatherall, 2001). Considering that multiple enzymatic activities could be simultaneously targeted for therapeutic purposes, the deubiquitinase BAP1 as well as additional epigenetic enzymatic activities that are regulated by BAP1 might provide novel targets for therapeutic intervention in inducing fetal globin synthesis for treatment of the β-globinopathies.

## Materials and Methods

### HUDEP-2 cell culture

HUDEP-2 cells were the kind gift of Dr. Yukio Nakamura (Kurita et al., 2013). HUDEP-2 cells were expanded in StemSpan SFEM medium (StemCell Technologies) supplemented with 50 ng/ml human stem cell factor (SCF) (Peprotech), 3 IU/ml erythropoietin (Amgen), 1 µg/ml doxycycline (Sigma), 1 µM Dexamethasone (Cayman) and 100 U/ml penicillin/streptomycin. To induce erythroid differentiation, HUDEP-2 cells were cultured in Iscove’s Modified Dulbecco’s Medium (IMDM) (Life Technologies) supplemented with 5% human AB serum (Sigma), 50 ng/ml SCF, 3 IU/ml erythropoietin (Amgen), 1 µg/ml Doxycycline (Sigma), 330 µg/ml holo-transferrin (Sigma), 10 µg/ml human insulin (Sigma), 2 IU/ml heparin (Sigma), and 100 U/ml penicillin/streptomycin.

### BioID

TR4-BirA*-expressing or parental wild type HUDEP-2 cells were incubated for 24h in StemSpan SFEM complete media supplemented with 50 µM biotin. After washing the cells with cold PBS, cell pellets (10^8^ cells) were resuspended in 4.5 mL of ice-cold RIPA buffer (50 mM Tris–HCl, pH 7.4, 150 mM NaCl, 1% NP-40, 1 mM EDTA, 1 mM EGTA, 0.1% SDS and 0.5% sodium deoxcycholate with 1 mM PMSF, 1 mM DTT and 1x Roche protease inhibitor cocktail). The cell lysates were sonicated twice on ice in a 15 ml conical tube for a total of 30s at (3s on/7s off, total pulse on 30s) with 40% amplitude (Fisher Scientific Sonic Dismembrator Model 500). 300 units of Benzonase (Sigma, E1014) were added and then rotated at 4°C for 60 minutes. Cell lysates were cleared by 16,500xg centrifugation at 4°C for 10 minutes. Supernatants were incubated with 250 µl Dynabeads MyOne Streptavidin C1 (ThermoFisher Scientific #65001; the beads were prewashed with RIPA buffer 3 times) in a 4°C room O/N. After incubation, the beads were washed with RIPA buffer 4 times, followed with 3 washes with fresh 50 mM NH_4_HCO_3_. The beads were frozen at −80 degrees until mass spectrometry identification.

### Protein identification by mass spectrometry and data analysis

The beads were resuspended in 50 µL of 0.1 M ammonium bicarbonate buffer (pH 8). Cysteines were reduced by adding 50 µL of 10 mM DTT and incubation at 45° C for 30’. Samples were cooled to room temperature and alkylation of cysteines was achieved by incubating with 65 mM 2-Chloroacetamide, in the dark, for 30’ at room temperature (RT). An overnight digestion with 1 µg of sequencing grade, modified trypsin was carried out at 37°C with constant shearing in a Thermomixer. Digestion was stopped by acidification and peptides were desalted using SepPak C18 cartridges following the manufacturer’s protocols (Waters). Samples were completely dried using a vacufuge, and the resulting peptides were dissolved in 8 µL of 0.1% formic acid/2% acetonitrile solution and 2 µL of the peptide solution was resolved on a nano-capillary reverse phase column (Acclaim PepMap C18, 2 micron, 50 cm, ThermoScientific) using a 0.1% formic acid/2% acetonitrile (Buffer A) and 0.1% formic acid/95% acetonitrile (Buffer B) gradient at 300 nl/min over a period of 180’ (2-22% buffer B for 110’, 22-40% for 25’, 40-90% for 5’ followed by holding in 90% buffer B for 5’ and requilibration with Buffer A for 35’). Eluent was directly introduced into Orbitrap Fusion tribrid mass spectrometer (Thermo Scientific, San Jose CA) using an EasySpray source. MS1 scans were acquired at 120K resolution (AGC target=1×10^6^; max IT=50 ms). Data-dependent collision induced dissociation MS/MS spectra were acquired using Top speed method (3 seconds) following each MS1 scan (NCE ∼32%; AGC target 1×10^5^; max IT 45 ms).

Proteins were identified by searching the MS/MS data against *Homo sapeiens* (Swissprot, v2016-11-30) using Proteome Discoverer (v2.1, Thermo Scientific). Search parameters included MS1 mass tolerance of 10 ppm and fragment tolerance of 0.2 Da; two missed cleavages were allowed; carbamidimethylation of cysteine was considered fixed modification and oxidation of methionine, deamidation of asparagine and glutamine were considered as potential modifications. False discovery rate (FDR) was determined using Percolator and proteins/peptides with a FDR of ≤1% were retained for final analysis.

### qPCR analysis of genomic DNA

Genomic DNA samples were prepared from mouse erythroid culture cells as described previously (Hosoya et al., 2013). Primers specific to NCoR1-floxed allele were used to quantify gene deletion by quantitative polymerase chain reaction (qPCR) with FastSybr Green Mastermix (Applied Biosystems) on an ABI Step One Plus (Applied Biosystems). Total allele of NCoR1 were amplified for normalization using primers specific to NCoR1 outside of loxP-franked region as described previously (Yu et al., 2014). All primer sequences used for qPCR analysis are listed in Supplemental Table 2.

### qRT-PCR analysis

Total RNA recovered from HUDEP-2 cell cultures was isolated using Trizol (ThermoFisher Scientific) according to the manufacturer’s instruction. cDNA was synthesized with SuperScript III Reverse Transcriptase (ThermoFisher Scientific). Total RNA from the murine *ex vivo* cultures was isolated using Trizol followed by the synthesis of complementary DNA (cDNA) using iScript cDNA synthesis kits (BioRad). qRT-PCR was performed using FastSybr Green Mastermix on an ABI Step One Plus. The abundance of human OAZ1 (Cui et al., 2015a; Cui et al., 2015b) and/or mouse GAPDH mRNA were used as the normalization controls. Sequences of all primers used for qRT-PCR are listed in Supplemental Table 2.

### ChIP assays

HUDEP-2 cells growing in the exponential phase were crosslinked by treatment with 1% formaldehyde at RT for 10 minutes with gentle shaking. Crosslinking was terminated by adding 0.125 M glycine for 5 min with gentle shaking at RT. Crosslinked cells were washed twice with cold PBS and 10^7^ cells were aliquoted for each ChIP assay. 10^7^ cells were resuspended in 1 ml Lysis buffer-1 [50 mM HEPES-KOH pH7.5; 140 mM NaCl; 1 mM EDTA; 10% glycerol; 0.5% NP-40; 0.25% Triton X-100; with 1 mM PMSF and 1x protease inhibitors (Sigma P8340)]. The cell suspension was rotated at 4°C for 10 min followed by 35 strokes in a type-B dounce homogenizer, on ice. The pellet was resuspended in 1 ml Lysis buffer-2 (10 mM Tris-Cl pH8; 200 mM NaCl; 1 mM EDTA; 0.5 mM EGTA; with 1 mM PMSF and 1× protease inhibitors) and rotated at RT for 10 minutes. The nuclei were then resuspended in 0.3 ml Lysis buffer-3 (10 mM Tris-Cl, pH8.0; 100 mM NaCl; 1 mM EDTA; 0.5 mM EGTA; 0.1% Na-deoxycholate; 0.5% N-lauroylsarcosine; with 1 mM PMSF and 1x protease inhibitors) in a 1.5 ml tube and sonicated. Sonication was performed at 40% amplitude with 10s pulse on, 20s pulse off, for 8 cycles (Fisher Scientific Sonic Dismembrator Model 500). After sonication, Triton X100 was added to 10% and cell debris was removed by centrifugation at 20,817 x g at 4°C for 10 minutes. 10% of the supernatant containing the chromatin was saved as the input control and the remaining 90% was brought to 500 µl by the addition of IP dilution buffer (50 mM Tris-HCl, pH7.4, 150 mM NaCl, 1 % NP-40, 0.25% deoxycholic acid, 1mM EDTA plus 1 mM PMSF and 1× protease inhibitors; Sigma P8340).

10 µl of protein A/G magnetic beads (ThermoFisher Scientific #88802) was washed two times with cold blocking solution (0.5% BSA in PBS), resuspended in 500 µl of blocking solution followed by rotation for 2 hr in a 4°C room. 2 µg of antibody or control rabbit IgG was added to the bead solution and then rotated O/N at 4°C. The beads were then washed one time with blocking solution and incubated with 10^7^ cell equivalents of chromatin for 12-16 hours at 4°C. The beads were gently washed with cold IP wash buffer (50 mM Tris-HCl, pH7.4, 150 mM NaCl, 1% NP-40, 0.25% deoxycholic acid, 1 mM EDTA) 3 times and 4°C PBS once. After washing, 300 µl of elution buffer (100 mM NaHCO_3_, 1% SDS) was used to remove the chromatin from the beads by incubation at 65°C with 900 rpm shaking for 2 hours. 36 µl of 5M NaCl was then added to reverse crosslink the chromatin, and the mixture incubated at 65°C O/N. After reverse crosslinking, DNase-free RNaseA digestion was performed at 37°C for 30 min followed by proteinase K treatment. The DNA sample was prepared by phenol extraction followed by column purification (Thermo Scientific) and subsequently analyzed by quantitative PCR. qPCR was performed using a FastSybr Green Mastermix (20 µl total reaction volume) on an ABI Step One Plus PCR machine with fast protocol settings [1 × 95°C, 20 sec; 40 × (95°C, 3 sec, 60°C, 30 sec), 1 × 95°C, 15 sec, 1 × 60°C for 60 sec with 0.3°C /sec increases up to 95°C for dissociation curve analysis]. Antibodies used for these experiments are listed in Supplemental Table 3.

### Yeast two-hybrid assays (Y2H)

Analysis of protein-protein interactions in yeast was performed as described previously (Vojtek and Hollenberg, 1995) using *Saccharomyces cerevisiae* L40 harboring *HIS3* and *lacZ* as reporter genes. All corepressor DNA sequences were amplified by PCR using mouse erythroleukemia (MEL) cDNA samples as templates. PCR fragments were subcloned into bait (pBTM116) or prey (pVP16) plasmids to generate LexA DNA binding domain (DBD) and VP16 activation domain (AD) fusion constructs, respectively. Point mutations were introduced into the constructs by site-directed mutagenesis. All constructs were verified by DNA sequencing. Interactions were tested in Y2H based on the induction of the *HIS3* reporter gene that allows yeast to grow in medium lacking histidine. Autoactivation of the *HIS3* reporter gene by bait plasmids was suppressed by addition of 3-aminotriazole (3-AT) into the selective medium. The expression of fusion proteins in yeast was confirmed by western blot using antibodies against LexA DBD or VP16 AD. An antibody recognizing phosphoglycerate kinase 1 was used as the loading control. Antibody information is shown in Supplemental Table 3.

### Lentiviral shRNAs

The human TR4 cDNA was clone into BioID plasmid (Addgene #74223) and the BirA*-TR4 fusion protein was then cut out and insert into CD550A-1 lentivirus plasmid (System Biosciences). The pLKO-puro lentiviral plasmids carrying shRNAs were obtained from Sigma-Aldrich. Human BAP1 shRNA clones TRCN0000007374 was used to knockdown BAP1 expression. TRC2 pLKO.5-puro non-mammalian shRNA were used as controls. Lentiviruses were prepared as previously described (Moffat et al., 2006). To knockdown BAP1, HUDEP-2 cells were transduced with lentiviruses carrying BAP1 or control (scrambled sequence) shRNAs by RetroNectin according to the product manufacturer (TaKaR #T100A/B). Transduced cells were selected with puromycin (0.25 μg/ml) 48 hr after transduction.

### *Ex vivo* fetal liver culture

The NCoR1flox/flox (Astapova et al., 2008) Mx1 Cre transgenic (Tg ^MX1Cre^) (Kuhn et al., 1995) mice were described previously. TER119 negative erythroblasts were purified from freshly isolated E14.5 fetal livers by a magnetic cell sorting system as previously described (Zhao et al., 2014), and expanded in StemPro-34 serum free media with nutrient supplement (Life Technologies) supplemented with 100 ng/ml mouse SCF (Peprotech), 1 μM Dexamethasone, 40 ng/ml mouse insulin-like growth factor 1 (IGF-1; R&D), 2 IU/ml erythropoietin, 100 U/ml penicillin/streptomycin, and 2 mM L-glutamine (Shearstone et al., 2011). To induce differentiation, cells were transferred to IMDM (L-glutamine, 25 mM HEPES) supplemented with 15% fetal bovine serum (Sigma), 1 % bovine serum albumin (StemCell Technologies), 10^−4^ M β-mercaptoethanol, 2 IU/ml erythropoietin, 200 μg/ml holo-transferrin (Sigma), 10 μg/ml human insulin (Sigma), and 100 U/ml penicillin/streptomycin (Zhao et al., 2014). To induce ID3/ID2 deletion *in vitro*, recombinant mouse interferon α (IFN-α; R&D) were added to culture during expansion phase at 1000 U/ml for 3 days.

### Co-immunoprecipitation (Co-IP)

Nuclear extracts (NE) were prepared as described (Folco EG et al. 2012. J Vis Exp). For each co-IP reaction, 200 µg of NE and 5 µg of antibody were mixed and brought to a final volume of 500 µl by the addition of sample buffer (10 mM HEPES-KOH, pH9.0, 1.5 mM MgCl_2_, 0.25 mM EDTA, 20% Glycerol, 0.3% NP-40, 0.5 mM DTT + 1 mM PMSF and 1× protease inhibitors (Sigma P8340)) and the mixture was rotated at 4°C O/N to promote antigen:antibody interaction. 25 µl of protein A/G magnetic beads (ThermoFisher Scientific #88802) were washed with PBS 3 times. After the final wash, the beads were blocked by treatment with Blocking buffer (10 mM HEPES-KOH, pH9.0, 1.5 mM MgCl_2_, 0.25 mM EDTA, 20% Glycerol, 200 µg/ml chicken egg albumin) and rotated O/N at 4°C. After blocking, the antigen:antibody mixture was incubated with the protein A/G beads at RT for 2 hours. The beads were then washed five times with Wash buffer (10 mM HEPES-KOH, pH9.0, 1.5 mM MgCl_2_, 0.25 mM EDTA, 20% Glycerol, 0.06% NP-40, 300 mM KCl + 1 mM PMSF and 1x protease inhibitors). Proteins were eluted from the beads by the addition of 1x SDS-PAGE loading buffer (Bio-Rad #161-0737) at 25°C for 10 minutes, electrophoresed, transferred to nylon membranes and then subjected to western blot detection. The antibodies used for Co-IP are listed in Supplemental Table 3.

### Western blotting

Western blotting was performed as previously described (Cui et al. 2011. MCB). The antibodies used for detection are listed in Supplemental Table 3.

### CRISPR mutagenesis

CRISPR-Cas9 targeted mutagenesis of NCoR1 was performed as described (Ran et al., 2013). The guide RNA (CCGGCAAATTGCCTCGGACA) was used to generate the NCoR1 mutant depicted in Figure 3A. The targeted (repairing) single strand oligonucleotide template sequence used is: 5’- ATAAAGGGCCTCCTCCAAAATCCAGATATGAGGAAGAGCTAAGGACCAGAGGG AAGACTACCATTACTGCAGCTAACTTCGCAGCCGCGGCCGCCACCCGACAGATAGCGTCCGATAAAGATGCGAGGGAACGTGGCTCTCAAAGTTCAGACTCTTCTAGTAGCTGTATGTATCTCAATCCGAGTTTCACAATGTGATGT-3’. The guide RNA was cloned in the px458 vector (Addgene #48138, with exchange of CMV to the EF1α promoter driving Cas9 expression). The vector and repairing template were transduced into HUDEP-2 cells by electroporation. EGFP positive cells were then sorted using a FACS Aria II, and single cell cloning was performed to generate the mutant cells.

### ChIP-seq analysis

The ChIP-seq data in K562 cells was obtained from ENCODE. NCoR1-1: ENCSR910JAI; NCoR1-2: ENCSR798ILC; eGFP-TR2: ENCSR178DEG; eGFP-TR4: ENCSR750LYM. The extent of peak overlap was calculated using bedtools (Quinlan and Hall, 2010). Gene Ontology (GO) enrichment was analyzed by ChIP-Enrich (Welch et al., 2014).

## Acknowledgements

We gratefully acknowledge the support, insights and comments of our partner colleagues [Yogen Saunthararajah (Cleveland Clinic), Joe DeSimone and Don Lavelle (U. Illinois-Chicago)] participating in a NHLBI Excellence in Hemoglobinopthies Research Award (U01 HL117658). We also appreciate numerous insightful comments from D. Lucas (Cincinnati Children’s Hospital) and the expert technical assistance of V. Basrur in the Department of Pathology Proteomics Resource Facility. We gratefully acknowledge support of shared facilities (sequencing and flow cytometry) from a NCI grant to the University of Michigan Comprehensive Cancer Center (P30 CA046592). Y.L. is supported by a fellowship from the Cooley’s Anemia Foundation, N.J. was supported by a Translational Scholars Award from the NHLBI (U01 HL117658) and M.P.L. was supported by a postdoctoral fellowship from the NIDDK (F32 DK108493).

## Author Contributions

Conceptualization: L.Y., N.J., K.C.L. and J.D.E.; resources, R.K. and Y.N.; formal Analysis, T.H. and G.M.; investigation, L.Y., N.J., M.P.L. and Q.W.; writing-original draft, L.Y., N.J. and J.D.E.; Supervision, A.B.V. and J.F.R.; funding acquisition, J.D.E.

## Supplemental Information

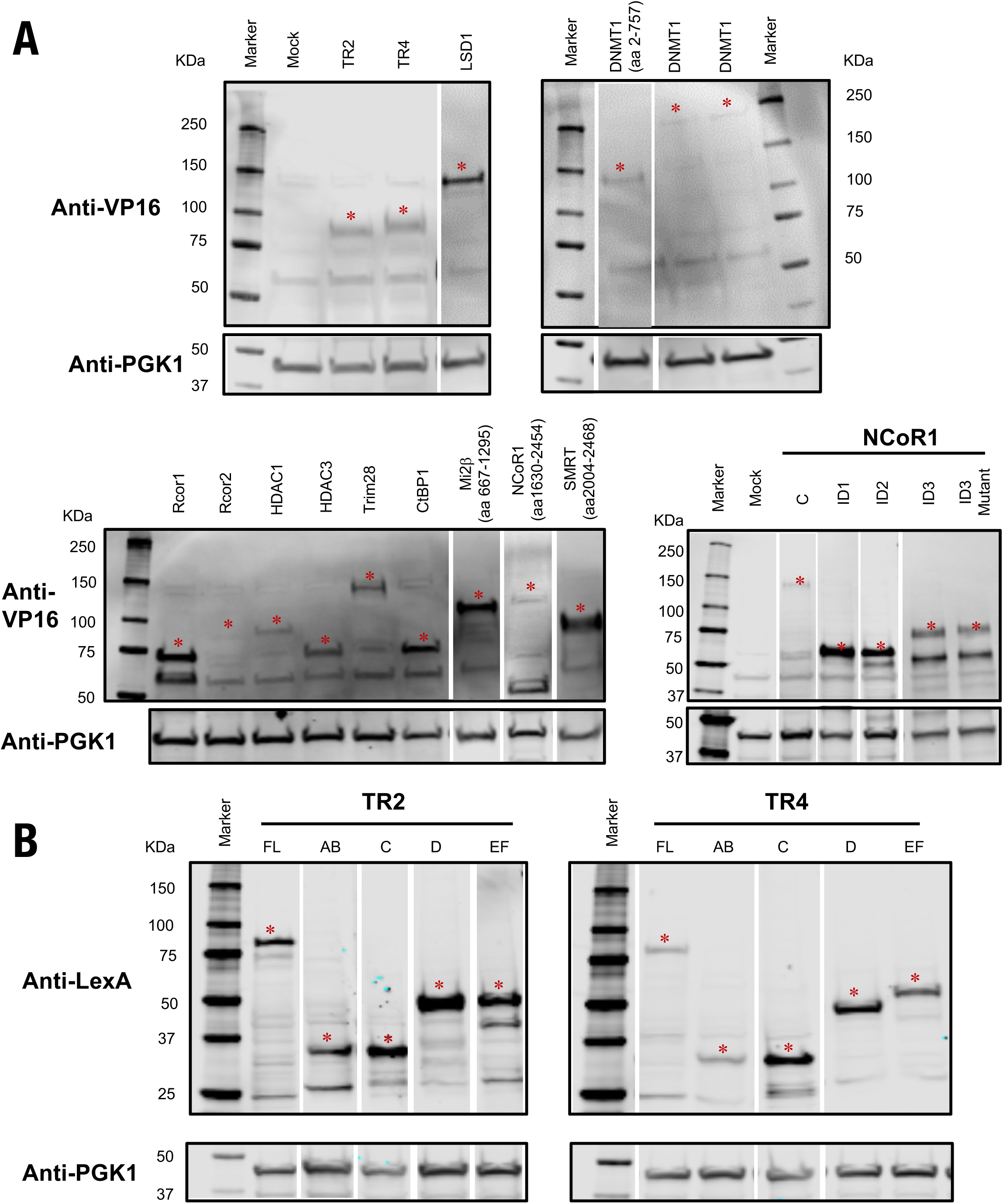
Expression of fusion proteins in *Saccharomyces cerevisiae* L40 for yeast two-hybrid experiments. Western blots were used to confirm the expression of (A) VP16 and (B) LexA fusion proteins in yeast using anti-VP16 activation domain or anti-LexA DNA binding domain antibodies, respectively. Expression of phosphoglycerate kinase 1 (PGK1) detected using an anti-PGK1 antibody was used as an internal control. * indicates the position of the expected fusion protein. White spaces between lanes were inserted to indicate the transition from non-continuous lanes or lanes spliced into the figure from different gels. FL, full length; aa, amino acid. Expression of LexA-NCoR1-RD1, LexA-NCoR1-RD2/3, Lex A-NCoR1-C and LexA-Sin3A are not shown.

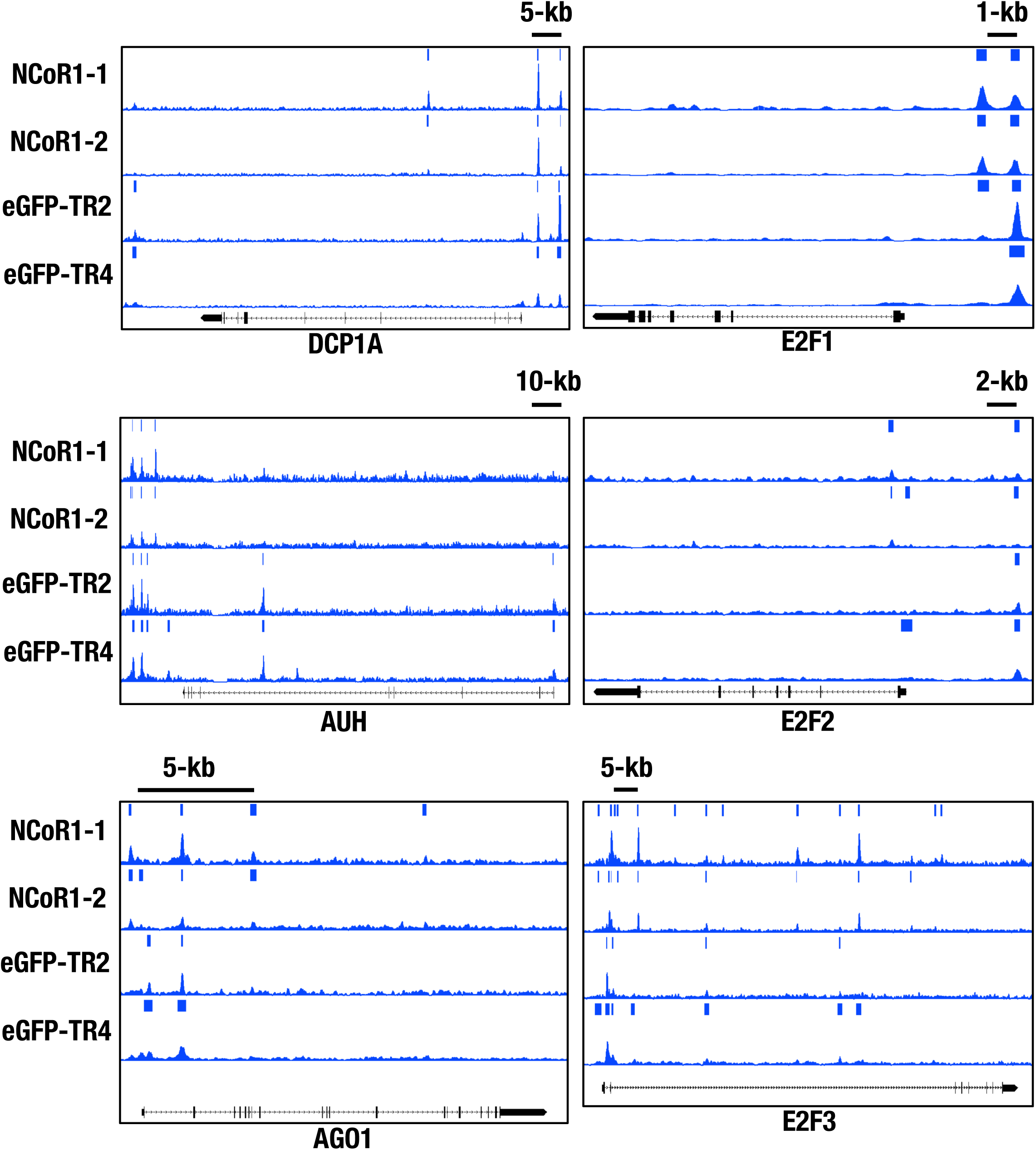
Representative TR2, TR4 and NcoR1 co-occupancy around the RNA catabolism-related genes DCP1A (mRNA-decapping enzyme 1A), AUH (AU RNA binding protein/enoyl-CoA hydratase) and AGO1 (Argonaute-1), and in the vicinity of the cell proliferation-related E2F transcription factor family genes.

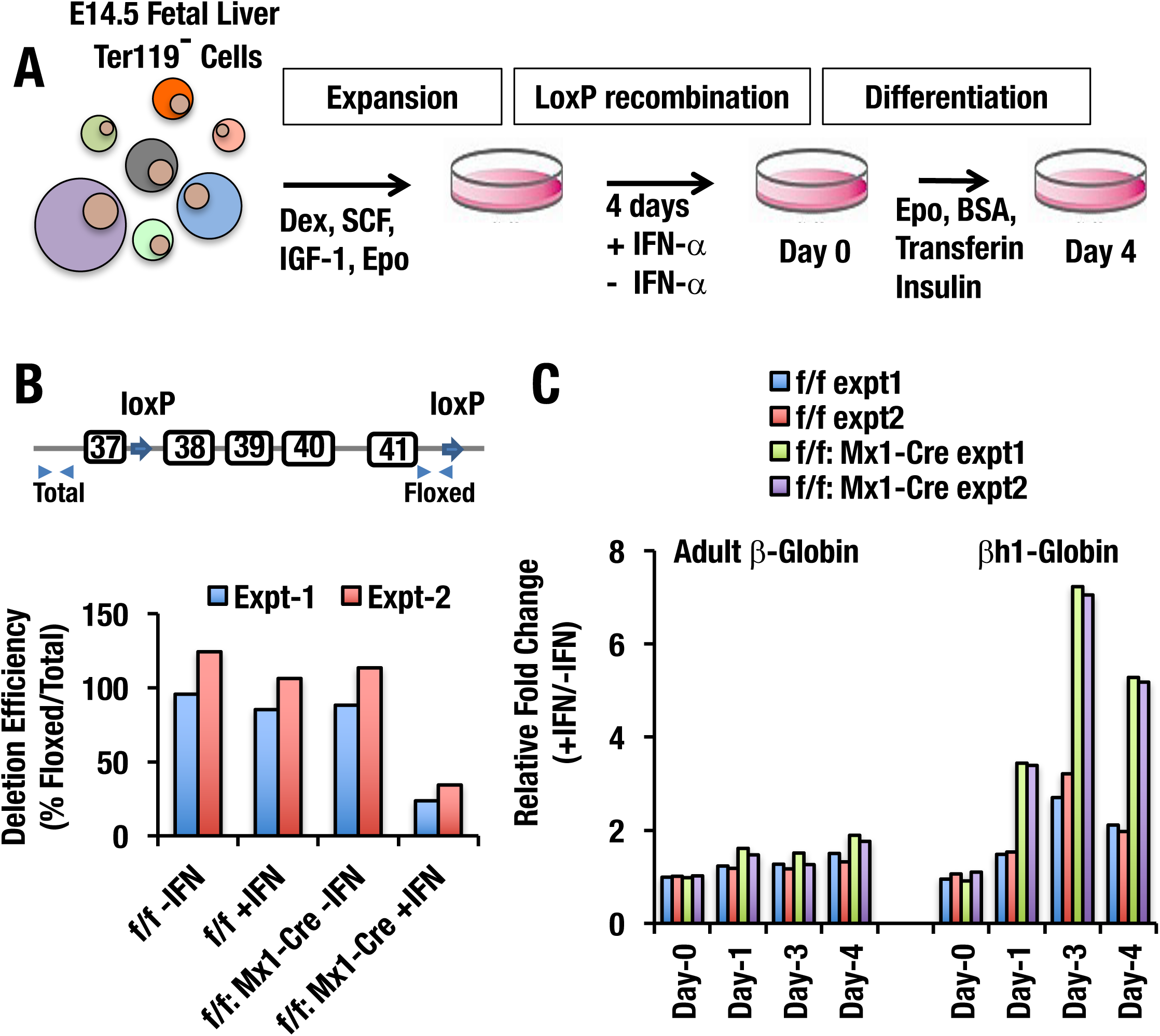
Conditional deletion of NcoR1 ID2/3 domain de-represses mouse *βh1-*globin gene expression. (A) Schematic diagram of *ex vivo* erythroid culture of E14.5 fetal liver Ter119- cells. (B) Conditional deletion efficiency of NcoR1 ID3 domain *ex vivo* after activation of Mx1-cre. Primers were designed to specifically detect unrearranged DNA at the loxP sites. (C) Loss of NcoR1 ID3 domain results in *βh1* embryonic murine globin gene induction *ex vivo*.

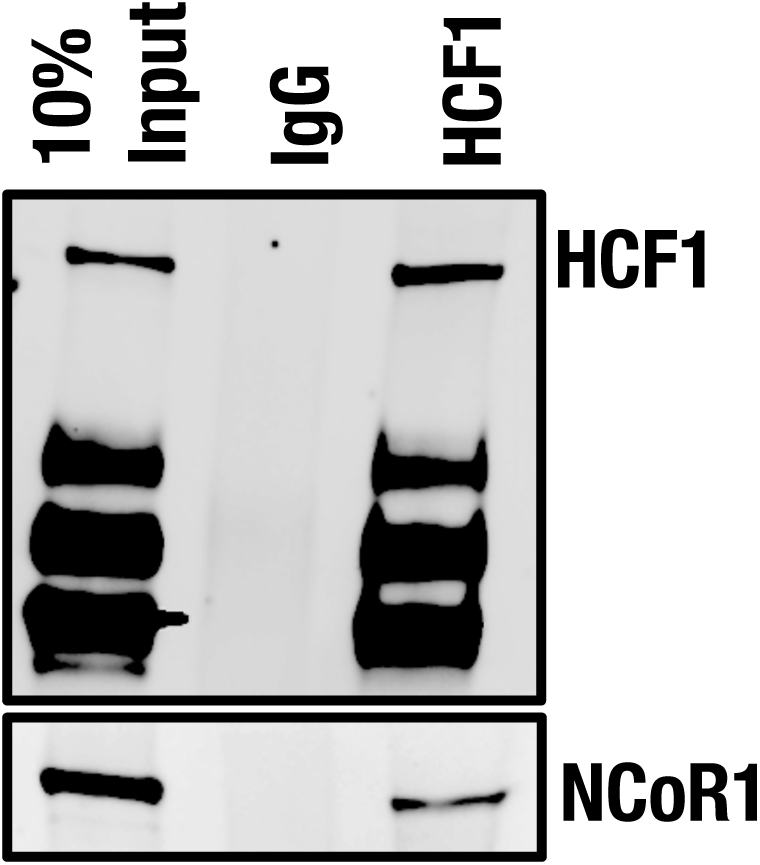
HCF1 and NCoR1 interaction is independent of TR4. Co-immune precipitation of HCF1 followed by western blot of HCF1 or NCoR1 in the NCoR1 CoRNR site mutant HuDEP-2 cell clone 1.

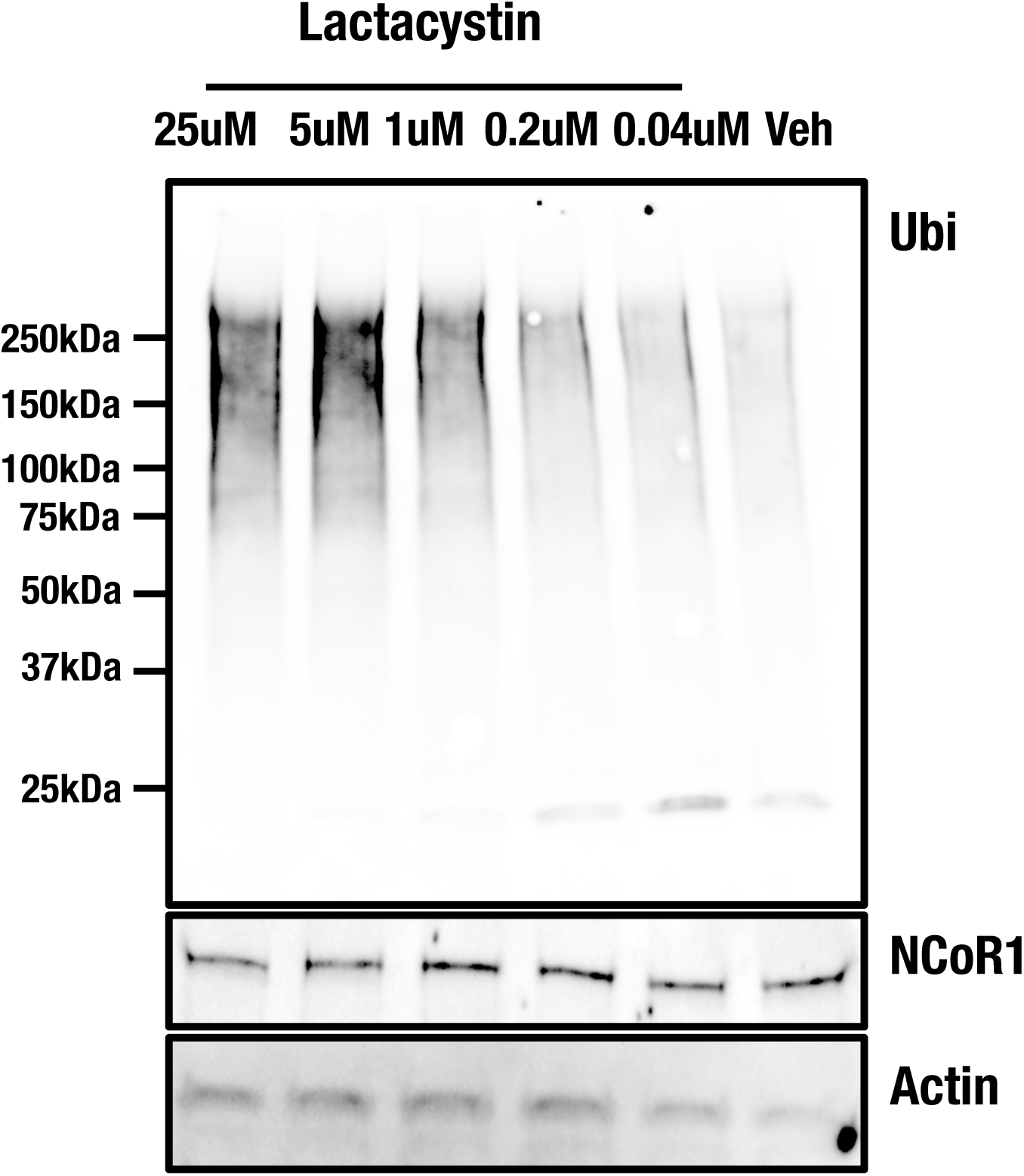
Proteasome inhibitor Lactacystin (Lac) treatment accumulates protein ubiquitination in HUDEP2 cells in dose-dependent way.

**Supplemental Table 1.** Proteins identified by BioID mass spectrometry. The total proteins identified from mass spectral analysis from control and TR4-BirA* HUDEP2 labeling experiments (3 each) is shown. Accession: UniprotKB protein accession number. Description: UniprotKB protein description. Sum PEP Score: protein score calculated as the negative log of the Posterior error probability (PEP) values of connected PSMs. Coverage: The percent calculated by dividing the number of amino acids in all found peptides by the total number of amino acids in the entire protein sequence. #Peptides: The number of distinct peptide sequences in the protein group. #PSMs: Peptide Spectrum Matches, the total number of identified peptide sequences for the proteins. #Unique Peptides: The number of unique peptide sequences identified of the proteins. #AAs: Number of amino acids of the proteins. MW [kDa]: Molecular weight of the <u>proteins. cale. pl:</u> calculated isoelectric point. Area: average of the peptide peak areas assigned to the protein. emPAI: Exponentially modified protein abundance index. Score: The protein score, which is the sum of the scores of the individual peptides calculated by Proteome Discoverer (v2.1, Thermo Scientific).

**Supplemental Table 2.**
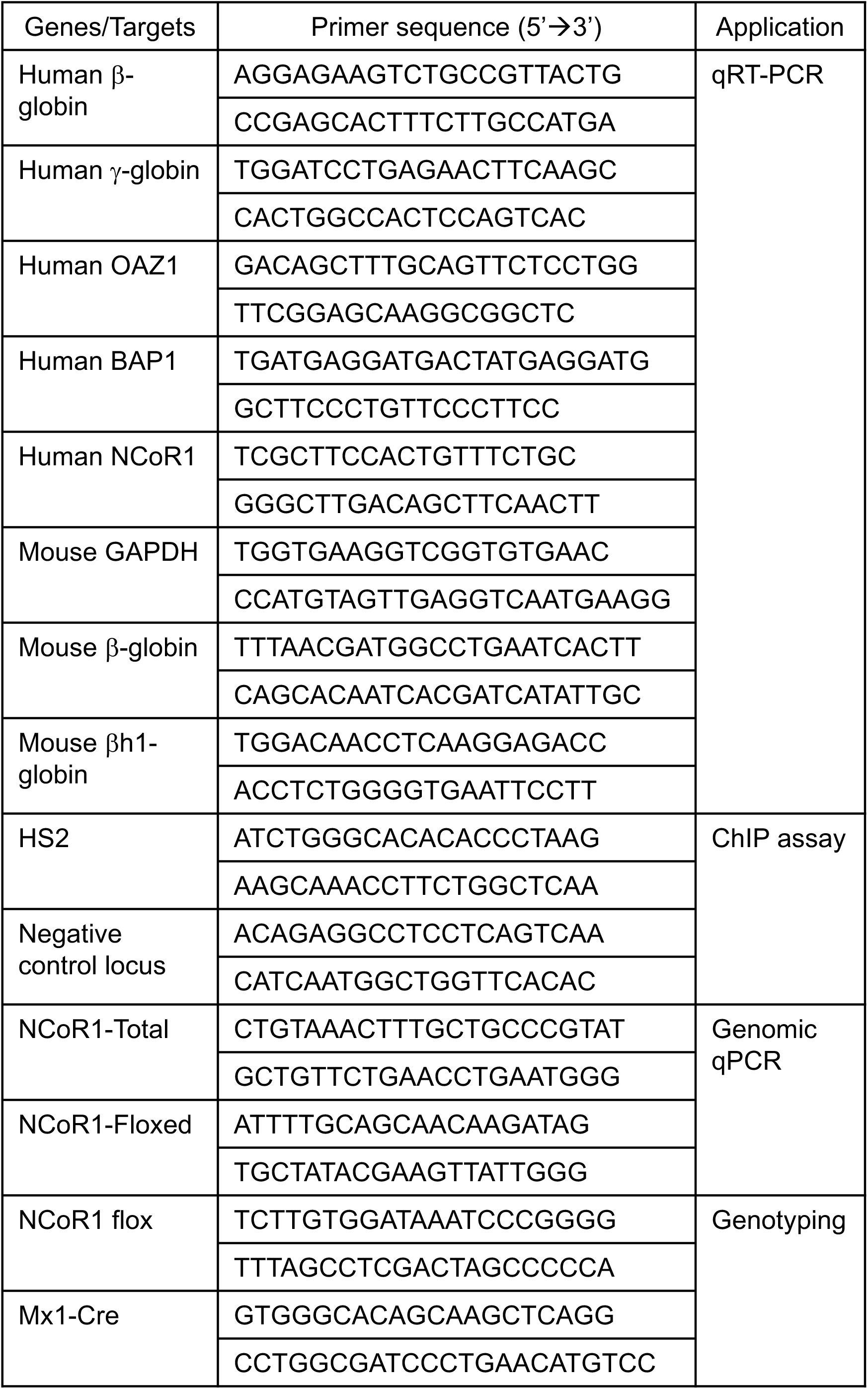
Primers used in this study.

**Supplemental Table 3.** Antibodies used in this study.

| Antibody<br>(Species) | Co-IP | WB | ChIP | Cat# |
| --- | --- | --- | --- | --- |
| NCoR1 (Rabbit) | + |  | + | A301-145A |
| HCF1 (Rabbit) | + |  |  | A301-399A |
| Ubi (Mouse) | + | + |  | sc-8017 |
| NCoR1 (Goat) |  | + |  | sc-1609 |
| TR4 (Rabbit) |  | + |  | Homemade |
| TR4 (Goat) |  | + |  | sc-8620 |
| LSD1 (Rabbit) |  | + | + | ab17721 |
| Dnmt1 (Mouse) |  | + |  | IMG-261A |
| HDAC1 (Rabbit) |  | + |  | ab7028 |
| HCF1 (Goat) |  | + |  | AF6254 |
| BAP1 (Mouse) |  | + |  | sc-28383 |
| LaminB (Goat) |  | + |  | sc-6217 |
| LexA DBD (Rabbit) |  | + |  | 06-719 |
| VP16 AD (Mouse) |  | + |  | sc-7545 |
| PGK1 (Mouse) |  | + |  | ab113687 |
| Streptavidin |  | + |  | 926-32230 |

## References

Astapova, I., Lee, L.J., Morales, C., Tauber, S., Bilban, M., and Hollenberg, A.N. (2008). The nuclear corepressor, NCoR, regulates thyroid hormone action in vivo. Proc Natl Acad Sci U S A 105, 19544–19549.

Bulger, M., and Groudine, M. (1999). Looping versus linking: toward a model for long-distance gene activation. Genes & development 13, 2465–2477.

Catic, A., Suh, C.Y., Hill, C.T., Daheron, L., Henkel, T., Orford, K.W., Dombkowski, D.M., Liu, T., Liu, X.S., and Scadden, D.T. (2013). Genome-wide map of nuclear protein degradation shows NCoR1 turnover as a key to mitochondrial gene regulation. Cell 155, 1380–1395.

Clegg, J.B., Weatherall, D.J., and Bodmer, W.F. (1983). 5-azacytidine for beta-thalassemia? Lancet 1, 536.

Cohen, R.N., Brzostek, S., Kim, B., Chorev, M., Wondisford, F.E., and Hollenberg, A.N. (2001). The specificity of interactions between nuclear hormone receptors and corepressors is mediated by distinct amino acid sequences within the interacting domains. Mol Endocrinol 15, 1049–1061.

Cui, S., Kolodziej, K.E., Obara, N., Amaral-Psarris, A., Demmers, J., Shi, L., Engel, J.D., Grosveld, F., Strouboulis, J., and Tanabe, O. (2011). Nuclear receptors TR2 and TR4 recruit multiple epigenetic transcriptional corepressors that associate specifically with the embryonic β-type globin promoters in differentiated adult erythroid cells. Mol Cell Biol 31, 3298–3311.

Cui, S., Lim, K.C., Shi, L., Lee, M., Jearawiriyapaisarn, N., Myers, G., Campbell, A., Harro, D., Iwase, S., Trievel, R.C., et al. (2015a). The LSD1 inhibitor RN-1 induces fetal hemoglobin synthesis and reduces disease pathology in sickle cell mice. Blood 126, 386–396.

Cui, S., Tanabe, O., Lim, K.C., Xu, H.E., Zhou, X.E., Lin, J.D., Shi, L., Schmidt, L., Campbell, A., Shimizu, R., et al. (2014). PGC-1 coactivator activity is required for murine erythropoiesis. Mol Cell Biol 34, 1956–1965.

Cui, S., Tanabe, O., Sierant, M., Shi, L., Campbell, A., Lim, K.C., and Engel, J.D. (2015b). Compound loss of function of nuclear receptors Tr2 and Tr4 leads to induction of murine embryonic beta-type globin genes. Blood 125, 1477–1487.

DeSimone, J., Heller, P., Hall, L., and Zwiers, D. (1982). 5-Azacytidine stimulates fetal hemoglobin synthesis in anemic baboons. Proc Natl Acad Sci U S A 79, 4428–4431.

Dey, A., Seshasayee, D., Noubade, R., French, D.M., Liu, J., Chaurushiya, M.S., Kirkpatrick, D.S., Pham, V.C., Lill, J.R., Bakalarski, C.E., et al. (2012). Loss of the tumor suppressor BAP1 causes myeloid transformation. Science 337, 1541–1546.

Engel, J.D., and Tanimoto, K. (2000). Looping, linking, and chromatin activity: new insights into beta-globin locus regulation. Cell 100, 499–502.

Folco, E.G., Lei, H., Hsu, J.L., and Reed, R. (2012). Small-scale nuclear extracts for functional assays of gene-expression machineries. J Vis Exp 64, pii: 4140.

Hosoya, T., Clifford, M., Losson, R., Tanabe, O., and Engel, J.D. (2013). TRIM28 is essential for erythroblast differentiation in the mouse. Blood 122, 3798–3807.

Hu, X., and Lazar, M.A. (1999). The CoRNR motif controls the recruitment of corepressors by nuclear hormone receptors. Nature 402, 93–96.

Jepsen, K., and Rosenfeld, M.G. (2002). Biological roles and mechanistic actions of co-repressor complexes. J Cell Sci 115, 689–698.

Kim, D.I., Jensen, S.C., Noble, K.A., Kc, B., Roux, K.H., Motamedchaboki, K., and Roux, K.J. (2016). An improved smaller biotin ligase for BioID proximity labeling. Molecular biology of the cell 27, 1188–1196.

Kuhn, R., Schwenk, F., Aguet, M., and Rajewsky, K. (1995). Inducible gene targeting in mice. Science 269,1427–1429.

Kurita, R., Suda, N., Sudo, K., Miharada, K., Hiroyama, T., Miyoshi, H., Tani, K., and Nakamura, Y. (2013). Establishment of immortalized human erythroid progenitor cell lines able to produce enucleated red blood cells. PLoS One 8, e59890.

Lambert, J.P., Tucholska, M., Go, C., Knight, J.D., and Gingras, A.C. (2015). Proximity biotinylation and affinity purification are complementary approaches for the interactome mapping of chromatin-associated protein complexes. Journal of proteomics 118, 81–94.

Lee, H.S., Lee, S.A., Hur, S.K., Seo, J.W., and Kwon, J. (2014). Stabilization and targeting of INO80 to replication forks by BAP1 during normal DNA synthesis. Nature communications 5, 5128.

Lee, M.P., Tanabe, O., LShi, L., Jearawiriyapaisarn, N., Lucas-Alcaraz, D., and Engel, J.D. (2017). The orphan nuclear receptor TR4 regulates erythroid cell proliferation and maturation. Blood, in press.

Ley, T.J., DeSimone, J., Noguchi, C.T., Turner, P.H., Schechter, A.N., Heller, P., and Nienhuis, A.W. (1983). 5-Azacytidine increases gamma-globin synthesis and reduces the proportion of dense cells in patients with sickle cell anemia. Blood 62, 370–380.

Lin, J.D. (2009). Minireview: the PGC-1 coactivator networks: chromatin-remodeling and mitochondrial energy metabolism. Mol Endocrinol 23, 2–10.

McCaffrey, P.G., Newsome, D.A., Fibach, E., Yoshida, M., and Su, M.S. (1997). Induction of gamma-globin by histone deacetylase inhibitors. Blood 90, 2075–2083.

Moffat, J., Grueneberg, D.A., Yang, X., Kim, S.Y., Kloepfer, A.M., Hinkle, G., Piqani, B., Eisenhaure, T.M., Luo, B., Grenier, J.K., et al. (2006). A lentiviral RNAi library for human and mouse genes applied to an arrayed viral high-content screen. Cell 124, 1283–1298.

Molokie, R., Lavelle, D., Gowhari, M., Pacini, M., Krauz, L., Hassan, J., Ibanez, V., Ruiz, M.A., Ng, K.P., Woost, P., et al. (2017). Oral tetrahydrouridine and decitabine for noncytotoxic epigenetic gene regulation in sickle cell disease: A randomized phase 1 study. PLoS medicine 14, e1002382.

Morales, J., Russell, J.E., and Liebhaber, S.A. (1997). Destabilization of human alpha-globin mRNA by translation anti-termination is controlled during erythroid differentiation and is paralleled by phased shortening of the poly(A) tail. J Biol Chem 272, 6607–6613.

Mottis, A., Mouchiroud, L., and Auwerx, J. (2013). Emerging roles of the corepressors NCoR1 and SMRT in homeostasis. Genes & development 27, 819–835.

Ngo, D.A., Aygun, B., Akinsheye, I., Hankins, J.S., Bhan, I., Luo, H.Y., Steinberg, M.H., and Chui, D.H. (2012). Fetal haemoglobin levels and haematological characteristics of compound heterozygotes for haemoglobin S and deletional hereditary persistence of fetal haemoglobin. Br J Haematol 156, 259–264.

Noguchi, C.T., Rodgers, G.P., Serjeant, G., and Schechter, A.N. (1988). Levels of fetal hemoglobin necessary for treatment of sickle cell disease. N Engl J Med 318, 96–99.

Palacios, I.M. (2013). Nonsense-mediated mRNA decay: from mechanistic insights to impacts on human health. Brief Funct Genomics 12, 25–36.

Perissi, V., Aggarwal, A., Glass, C.K., Rose, D.W., and Rosenfeld, M.G. (2004). A corepressor/coactivator exchange complex required for transcriptional activation by nuclear receptors and other regulated transcription factors. Cell 116, 511–526.

Perissi, V., Jepsen, K., Glass, C.K., and Rosenfeld, M.G. (2010). Deconstructing repression: evolving models of co-repressor action. Nat Rev Genet 11, 109–123.

Perissi, V., Scafoglio, C., Zhang, J., Ohgi, K.A., Rose, D.W., Glass, C.K., and Rosenfeld, M.G. (2008). TBL1 and TBLR1 phosphorylation on regulated gene promoters overcomes dual CtBP and NCoR/SMRT transcriptional repression checkpoints. Mol Cell 29, 755–766.

Perissi, V., Staszewski, L.M., McInerney, E.M., Kurokawa, R., Krones, A., Rose, D.W., Lambert, M.H., Milburn, M.V., Glass, C.K., and Rosenfeld, M.G. (1999). Molecular determinants of nuclear receptor-corepressor interaction. Genes & development 13, 3198–3208.

Qin, J., Zhou, Z., Chen, W., Wang, C., Zhang, H., Ge, G., Shao, M., You, D., Fan, Z., Xia, H., et al. (2015). BAP1 promotes breast cancer cell proliferation and metastasis by deubiquitinating KLF5. Nature communications 6, 8471.

Quinlan, A.R., and Hall, I.M. (2010). BEDTools: a flexible suite of utilities for comparing genomic features. Bioinformatics 26, 841–842.

Ran, F.A., Hsu, P.D., Wright, J., Agarwala, V., Scott, D.A., and Zhang, F. (2013). Genome engineering using the CRISPR-Cas9 system. Nat Protoc 8, 2281–2308.

Rivers, A., Vaitkus, K., Ibanez, V., Ruiz, M.A., Jagadeeswaran, R., Saunthararajah, Y., Cui, S., Engel, J.D., DeSimone, J., and Lavelle, D. (2016). The LSD1 inhibitor RN-1 recapitulates the fetal pattern of hemoglobin synthesis in baboons (P. anubis). Haematologica 101, 688–697.

Rivers, A., Vaitkus, K., Ruiz, M.A., Ibanez, V., Jagadeeswaran, R., Kouznetsova, T., DeSimone, J., and Lavelle, D. (2015). RN-1, a potent and selective lysine-specific demethylase 1 inhibitor, increases gamma-globin expression, F reticulocytes, and F cells in a sickle cell disease mouse model. Exp Hematol 43, 546–553 e541-543.

Roux, K.J., Kim, D.I., Raida, M., and Burke, B. (2012). A promiscuous biotin ligase fusion protein identifies proximal and interacting proteins in mammalian cells. J Cell Biol 196, 801–810.

Shearstone, J.R., Pop, R., Bock, C., Boyle, P., Meissner, A., and Socolovsky, M. (2011). Global DNA demethylation during mouse erythropoiesis in vivo. Science 334, 799–802.

Shi, L., Cui, S., Engel, J.D., and Tanabe, O. (2013). Lysine-specific demethylase 1 is a therapeutic target for fetal hemoglobin induction. Nature medicine 19, 291–294.

Suzuki, M., Yamamoto, M., and Engel, J.D. (2014). Fetal globin gene repressors as drug targets for molecular therapies to treat the beta-globinopathies. Mol Cell Biol 34, 3560–3569.

Tanabe, O., Katsuoka, F., Campbell, A.D., Song, W., Yamamoto, M., Tanimoto, K., and Engel, J.D. (2002). An embryonic/fetal beta-type globin gene repressor contains a nuclear receptor TR2/TR4 heterodimer. EMBO J 21, 3434–3442.

Tanabe, O., McPhee, D., Kobayashi, S., Shen, Y., Brandt, W., Jiang, X., Campbell, A.D., Chen, Y.T., Chang, C., Yamamoto, M., et al. (2007). Embryonic and fetal beta-globin gene repression by the orphan nuclear receptors, TR2 and TR4. EMBO J 26, 2295–2306.

Vojtek, A.B., and Hollenberg, S.M. (1995). Ras-Raf interaction: two-hybrid analysis. Methods Enzymol 255, 331–342.

Waggoner, S.A., and Liebhaber, S.A. (2003). Regulation of alpha-globin mRNA stability. Exp Biol Med (Maywood) 228,387–395.

Weatherall, D.J. (2001). The thalassemias. The molecular basis of blood diseases, 183–226.

Welch, R.P., Lee, C., Imbriano, P.M., Patil, S., Weymouth, T.E., Smith, R.A., Scott, L.J., and Sartor, M.A. (2014). ChIP-Enrich: gene set enrichment testing for ChIP-seq data. Nucleic acids research 42, e105.

Wood, W.G., Weatherall, D.J., and Clegg, J.B. (1976). Interaction of heterocellular hereditary persistence of foetal haemoglobin with beta thalassaemia and sickle cell anaemia. Nature 264, 247–249.

Xu, J., Bauer, D.E., Kerenyi, M.A., Vo, T.D., Hou, S., Hsu, Y.J., Yao, H., Trowbridge, J.J., Mandel, G., and Orkin, S.H. (2013). Corepressor-dependent silencing of fetal hemoglobin expression by BCL11A. Proc Natl Acad Sci U S A 110, 6518–6523.

Yu, L., Moriguchi, T., Souma, T., Takai, J., Satoh, H., Morito, N., Engel, J.D., and Yamamoto, M. (2014). GATA2 regulates body water homeostasis through maintaining aquaporin 2 expression in renal collecting ducts. Mol Cell Biol 34, 1929–1941.

Zarrizi, R., Menard, J.A., Belting, M., and Massoumi, R. (2014). Deubiquitination of gamma-tubulin by BAP1 prevents chromosome instability in breast cancer cells. Cancer research 74, 6499–6508.

Zhao, B., Mei, Y., Yang, J., and Ji, P. (2014). Mouse fetal liver culture system to dissect target gene functions at the early and late stages of terminal erythropoiesis. J Vis Exp, e51894.

Zhou, X.E., Suino-Powell, K.M., Xu, Y., Chan, C.W., Tanabe, O., Kruse, S.W., Reynolds, R., Engel, J.D., and Xu, H.E. (2011). The orphan nuclear receptor TR4 is a vitamin A-activated nuclear receptor. J Biol Chem 286, 2877–2885.

